# Transcription factor induction of vascular blood stem cell niches *in vivo*

**DOI:** 10.1101/2021.11.03.467105

**Authors:** Elliott J. Hagedorn, Julie R. Perlin, Rebecca J. Freeman, Samuel J. Wattrus, Tianxiao Han, Clara Mao, Ji Wook Kim, Inés Fernández-Maestre, Madeleine L. Daily, Christopher D’Amato, Michael J. Fairchild, Raquel Riquelme, Brian Li, Dana A.V.E. Ragoonanan, Khaliun Enkhbayar, Emily L. Henault, Helen G. Wang, Shelby E. Redfield, Samantha H. Collins, Asher Lichtig, Song Yang, Yi Zhou, Balvir Kunar, Jesus Maria Gomez-Salinero, Thanh T. Dinh, Junliang Pan, Karoline Holler, Henry A. Feldman, Eugene C. Butcher, Alexander van Oudenaarden, Shahin Rafii, J. Philipp Junker, Leonard I. Zon

## Abstract

The hematopoietic niche is a supportive microenvironment comprised of distinct cell types, including specialized vascular endothelial cells that directly interact with hematopoietic stem and progenitor cells (HSPCs). The molecular factors that specify niche endothelial cells and orchestrate HSPC homeostasis remain largely unknown. Using multi-dimensional gene expression and chromatin accessibility analyses, we define a conserved gene expression signature and *cis*-regulatory landscape unique to sinusoidal endothelial cells in the HSPC niche. Using enhancer mutagenesis and transcription factor overexpression, we elucidate a transcriptional code involving members of the Ets, Sox and Nuclear Hormone Receptor families that is sufficient to induce ectopic niche endothelial cells that associate with mesenchymal stromal cells and support the recruitment, maintenance and division of HSPCs *in vivo*. These studies set forth an approach for generating synthetic HSPC niches, *in vitro* or *in vivo*, and for effective therapies to modulate the endogenous niche.

---

Hematopoietic stem and progenitor cells (HSPCs) are a rare population of cells capable of reconstituting the entire blood system^1^. In bone marrow, multiple cell types comprise the HSPC niche - primarily endothelial cells (ECs) and perivascular mesenchymal stromal cells^2–7^. Distinct endothelial subtypes differentially regulate HSPCs: arterial ECs (AECs) promote HSPC quiescence, while sinusoidal ECs (SECs) support HSPC differentiation and mobilization^8–10^. Specialized bone marrow ECs play a critical role in niche reconstruction and hematopoietic recovery after myelosuppression^11,12^, and ECs support HSPCs outside the bone marrow during development and stress-induced hematopoiesis^13^.

HSPCs are born in the aorta-gonad-mesonephros region and then migrate to a transient fetal niche, the liver in mammals or a venous plexus in the tail of fish called the caudal hematopoietic tissue (CHT)^1,14^. HSPCs expand in these sites for several days before migrating to the adult niche, the bone marrow in mammals or kidney marrow in fish. The CHT is comprised primarily of low-flow venous SECs surrounded by mesenchymal stromal cells^14–19^. As HSPCs lodge in the CHT, ECs reorganize to form supportive pockets, which together with stromal cells form a niche^17^. Specific signaling molecules, adhesion proteins and transcription factors are implicated in mediating cross-talk and physical interaction between stem cells and ECs in the niche^2,20–26^. Understanding the transcriptional regulation of these molecules could guide new strategies to improve the efficacy and availability of bone marrow transplantation.

## Results

### An endothelial gene expression signature unique to HSPC niches

To investigate gene expression in the CHT, we performed RNA tomography (tomo-seq)^27^ on the zebrafish tail at 72 hours post fertilization (hpf; Fig. 1a). This revealed clusters of gene expression corresponding to specific tissues along the dorsal-ventral axis, including spinal cord, notochord, muscle, epidermis and hematopoietic populations (Fig. 1b and Extended Data Fig. 1a). 144 genes were enriched in the CHT (Fig. 1b and Supplementary Table 1). Using EC-specific RNA-seq, published myeloid RNA-seq datasets^28^ and whole mount *in situ* hybridization (WISH), we identified 29/144 genes that were selectively enriched in CHT ECs (Fig. 1b and c, Extended Data Fig. 1b and Supplementary Table 2). Using published whole kidney and new EC-specific single cell RNA-seq data, we found that 23 of these 29 genes were expressed by venous SECs in the adult kidney (Fig. 1d and Extended Data Fig. 1c), a population associated with hematopoiesis in fish^29^. The orthologs for 21/29 CHT EC genes were enriched in the ECs of mammalian hematopoietic organs^30^ – the fetal liver and/or adult bone marrow, at stages when these tissues support hematopoiesis (Fig. 1e). Thus, the niche endothelial signature identified in the CHT is largely conserved across species and hematopoietic development.

**Figure 1.**
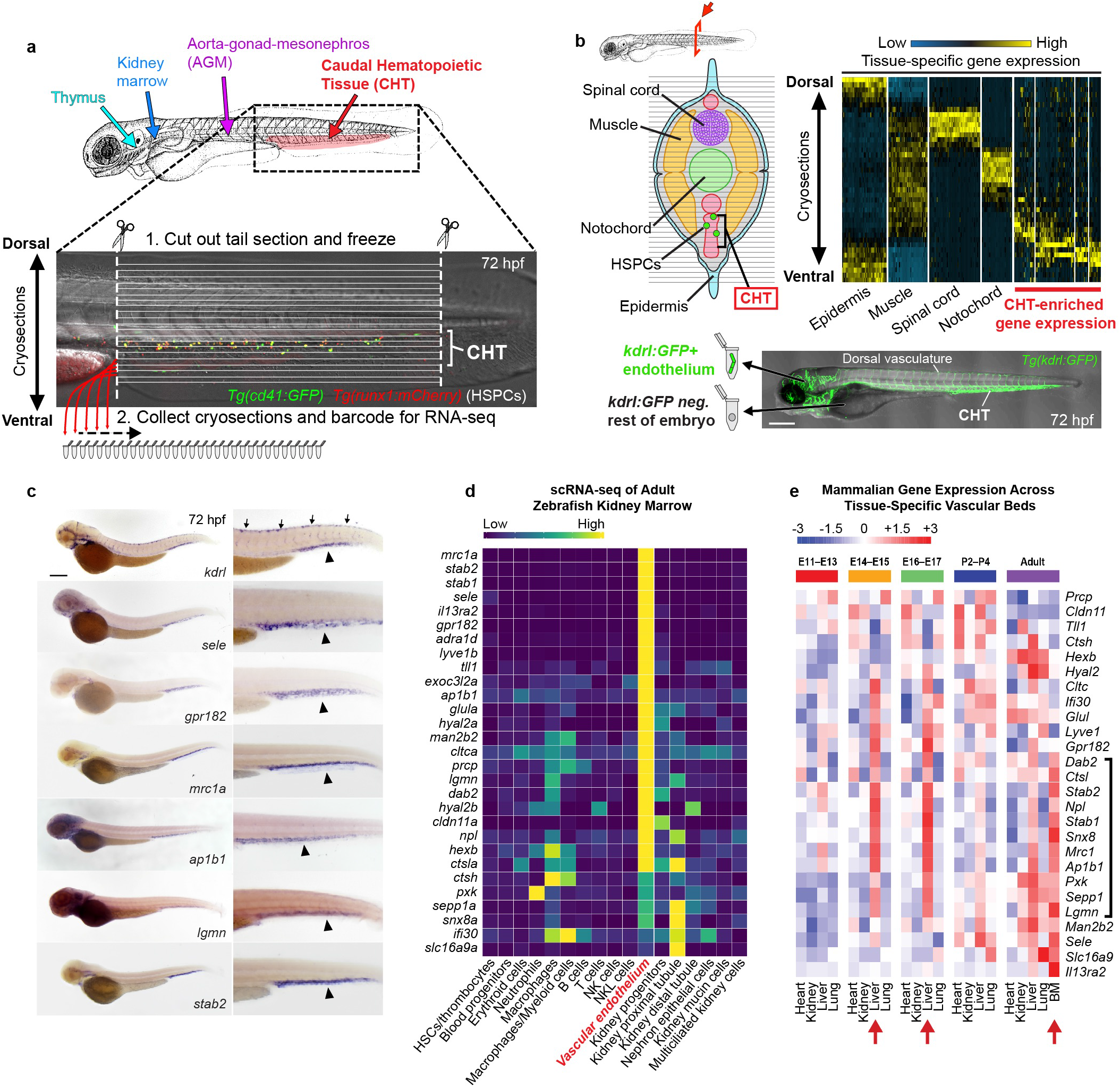
An endothelial gene expression signature unique to HSPC niches. **a,** Schematic diagram illustrates the haematopoietic tissues of the zebrafish embryo (top) and the sectioning strategy used to perform RNA tomography (tomo-seq) on the CHT (bottom; double transgenic embryo carrying the HSPC markers *cd41:GFP* and *runx1:mCherry* is shown). **b**, Schematic cross-section (upper left) and hierarchical clustering heat map (upper right) reveal clusters of gene expression that correspond to distinct tissues along the dorsal-ventral axis of the zebrafish tail. Schematic (bottom) depicts strategy using *kdrl:GFP* transgenic embryos and FACS to isolate ECs from whole embryos for analysis by RNA-seq. **c**, Images show whole mount *in situ* hybridization (WISH) for the pan-endothelial gene *kdrl* (top panel) and CHT EC-enriched genes identified by tomo-seq and tissue-specific RNA-seq (bottom panels). Arrows point to expression in dorsal vasculature and arrowheads point to expression in the CHT. **d**, Heat map shows the expression of the 29 CHT EC genes in the different cell populations that comprise the adult zebrafish kidney marrow. Spectral scale reports normalized expression. **e**, Heat map shows the expression of orthologs of the zebrafish CHT EC genes in ECs from different organs of the mouse at different stages of development and postnatal transition to adulthood. Red arrows denote haematopoietic tissues at the respective stage of development. Black bracket denotes genes enriched in fetal liver ECs at the E14-17 stages and then later in the adult bone marrow. Spectral scales report z-scores. BM: Bone Marrow. Scale bars represent 250 μm in this and all subsequent figures unless noted otherwise.

### Endothelial niche-specific cis-regulatory elements

To isolate CHT ECs, we generated GFP reporter transgenes using 1.3 or 5.3 kb upstream regulatory sequences for two highly expressed CHT endothelial genes known to promote hematopoietic cell adhesion: *mrc1a* and *sele^25,31,32^*. We crossed these reporters to the pan-endothelial marker *kdrl:mCherry*. For both the *mrc1a 1.3kb:GFP* and *sele 5.3kb:GFP* transgenes, the highest vascular expression was observed in venous SECs of the CHT, which directly interact with HSPCs and *cxcl12a:DsRed2*+ stromal cells (Extended Data Fig. 2a-e). Selective GFP expression was similarly observed in kidney marrow ECs (Extended Data Fig. 2f-g), consistent with these transgenes being markers of niche ECs.

To investigate transcriptional control of niche-specific gene expression, we dissociated double positive *mrc1a 1.3kb:GFP*; *kdrl:mCherry* embryos and isolated four populations for RNA-seq and ATAC-seq: GFP^+^; mCherry^+^ (CHT ECs), GFP^-^; mCherry^+^ (non-CHT ECs), GFP^+^; mCherry^-^ (mesenchymal cells in the tail fin), and GFP^-^; mCherry^-^ (negative remainder of the embryo; Fig. 2a). We identified 6,848 regions of chromatin uniquely open in CHT ECs (Supplementary Table 3). Of the 29 CHT EC genes, 26 had an ATAC-seq element within 100 kb of the transcriptional start site accessible only in CHT ECs (Fig. 2b, Extended Data Fig. 3a and Supplementary Table 2). Similar regions of chromatin accessibility were detected when using the *sele 5.3kb:GFP* transgene (Extended Data Fig. 3d and Supplementary Table 3). To test whether the uniquely accessible regions of chromatin are tissue-specific enhancers, we cloned 15 of the elements, fused them to a minimal promoter and GFP, and injected them into zebrafish embryos. 12/15 constructs showed GFP enrichment in CHT ECs at 60-72 hpf (Fig. 2b, Extended Data Fig. 3a and Supplementary Table 4). Conversely, 6/6 pan-endothelial ATAC-seq elements drove mosaic GFP expression in ECs throughout the entire embryo (Extended Data Fig. 3b), illustrating the specificity of the CHT elements. Stable integration of the enhancer transgenes confirmed the expression observed in F0 animals (Extended Data Fig. 3c).

**Figure 2.**
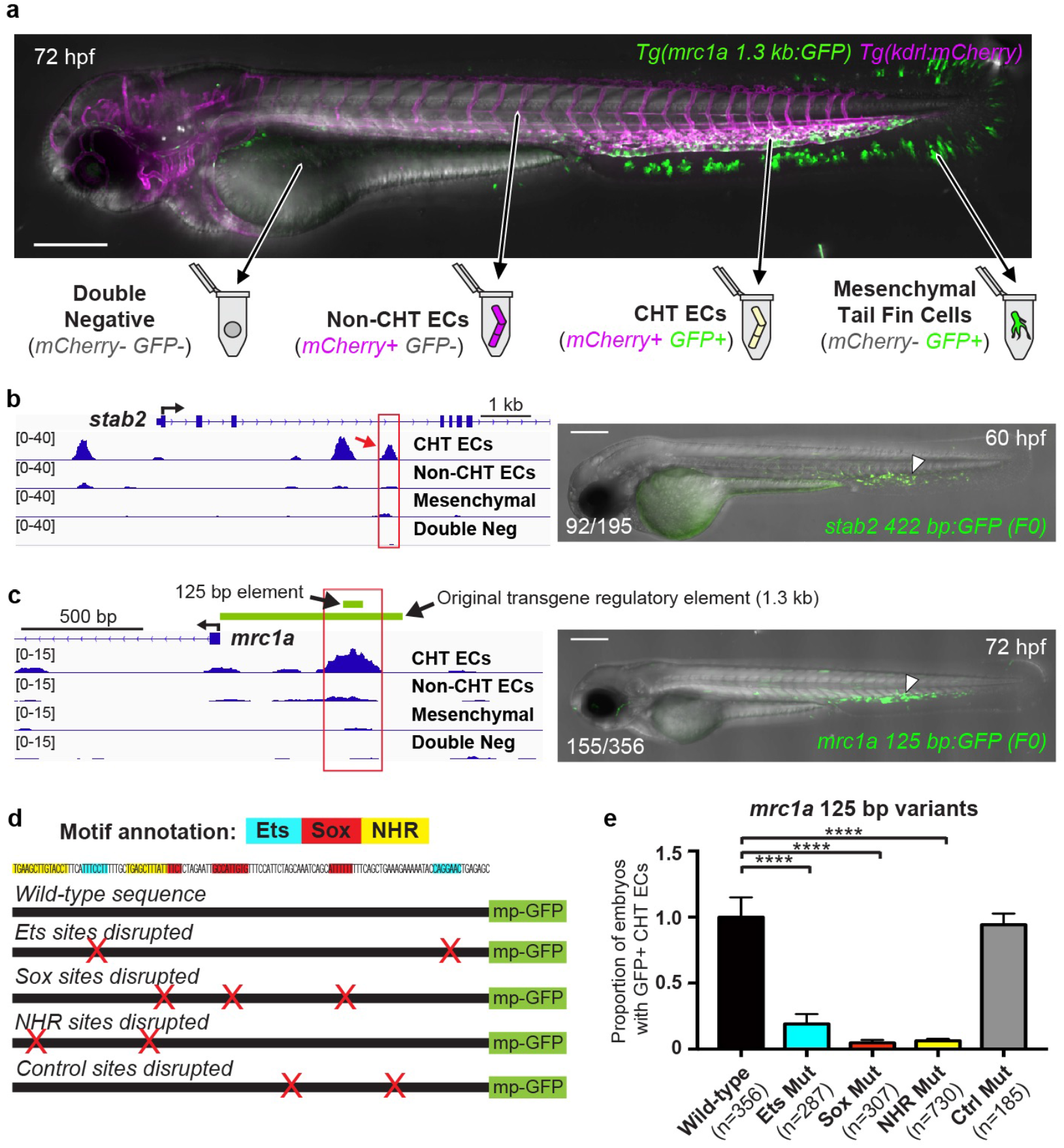
Endothelial niche-specific cis-regulatory elements. **a**, Image and schematic depict the four cell populations that were isolated from *mrc1a 1.3kb:GFP*^+^; *kdrl:mCherry*^+^ double positive embryos for analysis by ATAC-seq. **b**, Gene tracks show regions of chromatin that were uniquely open in the GFP^+^mCherry^+^ CHT EC fraction. Image on the right shows an embryo injected with a CHT EC enhancer-GFP reporter construct corresponding to the red boxed region (red arrow). Arrowhead points to GFP expression in CHT ECs. **c**, Gene tracks show a region of chromatin (red box) upstream of *mrc1a* that is uniquely open in the double positive CHT EC fraction but not the other three cell populations. Green bars denote the position of the 125 bp enhancer sequence and the 1.3 kb sequence used to generate the reporter transgenes. Image on the right shows transient GFP expression in an F0 embryo injected with the 125 bp enhancer sequence coupled to a minimal promoter and GFP. **d**, Wild-type sequence of the 125 bp *mrc1a* enhancer is shown, annotated with colors highlighting the Ets, Sox and NHR binding motifs. Schematic depicts enhancer-reporter constructs in which each class of motif or control regions was targeted by mutation. Red X’s denote the location of targeted sites. mp-GFP: mouse *Beta-globin* minimal promoter fused to GFP. **e**, Graphs report the frequency of embryos with GFP expression in CHT ECs after injection with wild-type sequences or mutated variants of the *mrc1a* 125 bp enhancer. Data is normalized to the wild-type control (44% (155/356)). Mean +/− s.e.m., One-way ANOVA with Dunnett’s multiple comparisons test; ***P<0.001, ****P<0.0001.

To determine a minimal sequence sufficient to drive CHT EC gene expression, we cloned 125 bp and 158 bp sequences from the strongest ATAC-seq signal upstream of *mrc1a* and *sele*, respectively (Fig. 2c and Extended Data Fig. 4a). When coupled to a minimal promoter, these elements drove GFP expression that was selectively enriched in CHT ECs in 44% (125 bp, *mrc1a*; 155/356) and 23% (158 bp, *sele*; 176/775) of embryos (Fig. 2c and Extended Data Fig. 4a-c). Upon stable integration of each transgene, GFP expression was restricted to CHT ECs (Extended Data Fig. 4d). Single cell RNA-seq of FACS-purified ECs from *mrc1a 125bp:GFP^+^; kdrl:mCherry^+^* embryos confirmed that GFP^+^ cells selectively expressed the 29-gene niche endothelial signature (Extended Data Fig. 5a). Transcripts for GFP and some of the 29 genes were also detected in a population of head lymphatic ECs (Extended Data Fig. 5a), however, a direct comparison between the CHT EC and head lymphatic EC populations revealed substantial differences in gene expression, including the *bona-fide* vascular niche factors *vcam1b, cxcl12a* and *sele*, which were expressed by CHT ECs but not head lymphatic ECs (Extended Data Fig. 5a and Supplementary Table 5), consistent with the inability of the head lymphatic ECs to recruit and support HSPCs. Within the CHT, *mrc1a 125bp:GFP* expression turned on shortly before HSPC colonization, increased in intensity through 8 dpf (coincident with HSPC expansion), and then decreased steadily as HSPCs exited the CHT (Extended Data Fig. 5b and Supplementary Video 1). A similar dynamic was observed in the kidney marrow, where GFP expression was observed shortly before HSPC colonization (Extended Data Fig. 5b), consistent with a role for *mrc1a* in promoting adhesive interactions between HSPCs and the vascular niche.

To identify transcription factors that bind CHT EC enhancers, we performed motif enrichment analysis of the 6,848 regions of chromatin uniquely accessible in CHT ECs. This revealed that Ets, SoxF and Nuclear Hormone Receptor (NR2F2/RORA/RXRA, specifically, abbreviated hereafter as NHR) binding motifs were highly enriched (Extended Data Fig. 3e). In contrast, 4,522 pan-endothelial elements were enriched for Ets, but not SoxF or NHR binding motifs (Supplementary Table 3; Extended Data Fig. 3e). To test whether the Ets, SoxF and NHR sites were required for expression, we generated variants of the 125 bp and 158 bp enhancer sequences with each motif class mutated (Fig. 2d and Extended Data Fig. 4e). In each case, a significant reduction or complete loss of GFP expression in CHT ECs was observed (Fig. 2e and Extended Data Fig. 4e). GFP expression was unperturbed in embryos injected with control constructs with mutations between the Ets, SoxF and NHR motifs (Fig. 2d and e, and Extended Data Fig. 4e). Electrophoretic mobility shift assays demonstrated that NR2F2 (also known as COUP-TFII), a NHR that promotes venous identity^33^, was able to bind the NHR motifs in the *mrc1a* 125 bp and *sele* 158 bp enhancers (Extended Data Fig. 4f). Together, this work defines a *cis*-regulatory landscape unique to niche ECs and suggests that Ets, Sox and NHR transcription factors drive niche endothelial development.

### Defined factors induce niche endothelial expression

To determine which transcription factors might bind the Ets, Sox and NHR motifs *in vivo*, we examined our bulk RNA-seq data from the double positive CHT ECs. The most highly expressed factors were *fli1a*, *etv2*, *ets1*, *sox18*, *sox7*, *nr2f2* and *rxraa* (Supplementary Table 6). To test whether these factors were sufficient to induce ectopic niche endothelial gene expression, we injected a pool of constructs encoding orthologs for the seven factors driven by a ubiquitous *(ubĩ)* promoter into zebrafish embryos and examined *mrc1a* and *sele* expression by WISH at 60-72 hpf (Fig. 3a). Strikingly, 17% (12/69) of the 7-factor-injected embryos had ectopic *mrc1a* expression in the head, trunk and over the yolk, whereas controls did not (0/56; Extended Data Fig. 6a and c). Similar results were obtained with WISH for *sele* or when factors were injected into *mrc1a 1.3kb:GFP* and *sele 5.3kb:GFP* embryos (Fig. 3b and Extended Data Fig. 6a-b and d).

**Figure 3.**
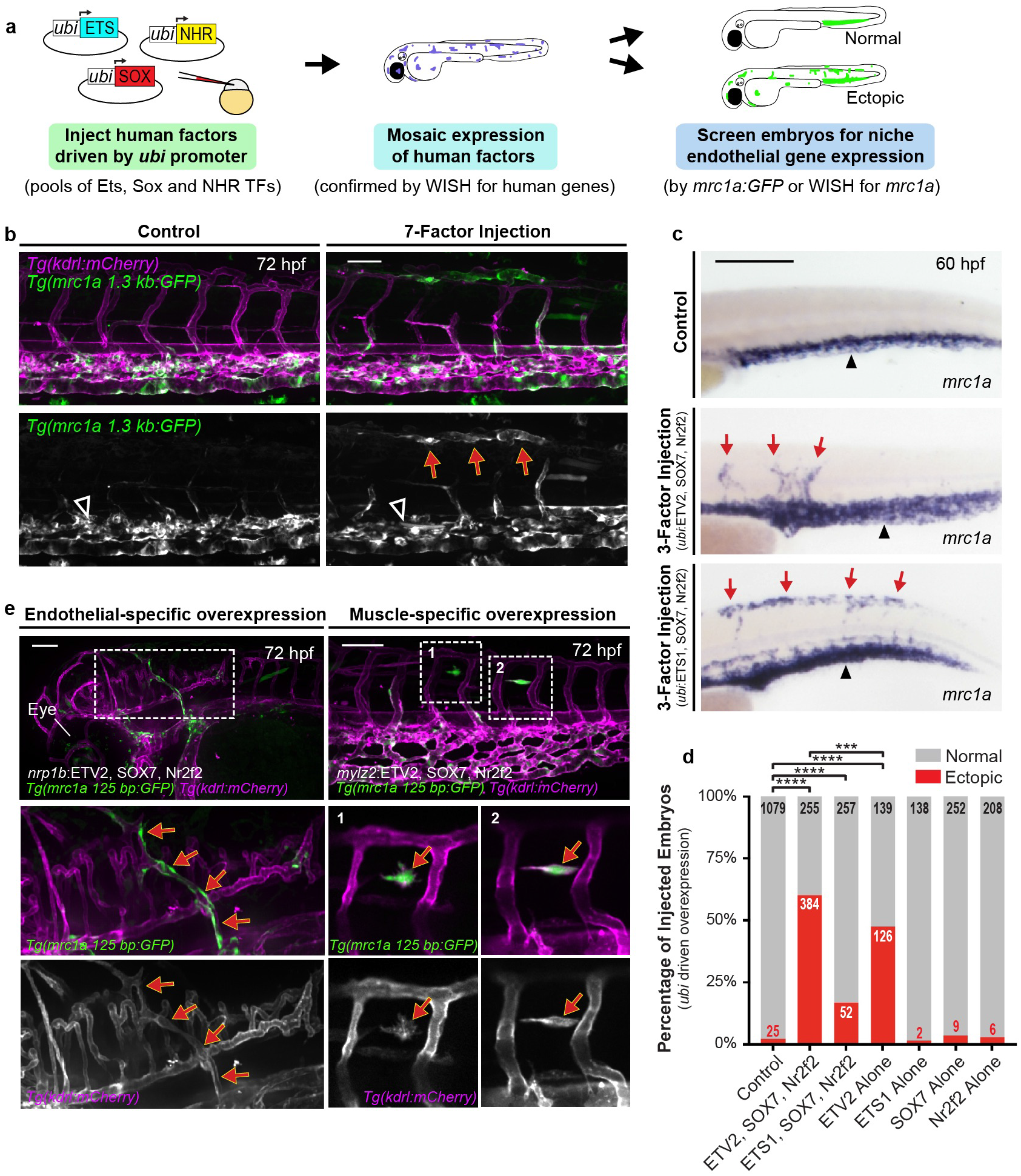
Defined factors induce niche endothelial expression. **a**, Schematic depicts the strategy used in transcription factor overexpression experiments. **b**, Images show *mrc1a 1.3kb:GFP; kdrl:mCherry* double transgenic embryos that were injected with control DNA (left) or a pool of seven transcription factors (right) from the Ets, Sox and NHR families (FLI1, ETV2, ETS1, SOX7, Sox18, Nr2f2 and RXRA). Red arrows denote regions of ectopic expression and black arrowheads point to normal domains of expression in all panels of this figure. **c**, Images show WISH for *mrc1a* in a control embryo (top) or after injection of a 3-factor pool containing ETV2, SOX7 and Nr2f2 (middle) or ETS1, SOX7 and Nr2f2 (bottom). **d**, Graph reports quantification of the percentage of injected embryos that displayed ectopic *mrc1a* WISH staining after transcription factor overexpression. Chi Square Test and Fisher’s exact test for pairwise comparisons with the Holm step-down process to correct for multiple comparisons; ***P<0.001, ****P<0.0001. **e**, Images show ectopic expression in *mrc1a 125bp:GFP*; *kdrl:mCherry* double positive embryos that were injected with endothelial-specific *nrp1b:ETV2*, SOX7, Nr2f2 (left) or muscle-specific *mylz2:ETV2*, SOX7, Nr2f2 (right) plasmids. Magnification of boxed regions is shown at the bottom. Scale bars represent 100 μm in **b** and **e**, and 250 μm in **c**.

Our mutant enhancer experiments indicated that at least one factor from each of the three families was required for niche EC gene expression, which led us to ask whether a combination of just three factors, which one from each family, might be sufficient to induce ectopic niche EC gene expression. ETV2 is a pioneer factor essential for specification of early mesodermal progenitors into vascular cell fates^34,35^. Forced expression of ETV2 in nonvascular cells induces reprogramming towards an early endothelial fate that can generate many types of vasculature^36–38^. Previous work in zebrafish has shown the importance of SoxF factors *(sox7* and *sox18)* and *nr2f2* during arterial-venous specification^39^. We therefore hypothesized that a combination of three of these factors – ETV2, SOX7 and Nr2f2 – might be sufficient to induce ectopic niche endothelial gene expression. We injected a pool of *ubi*-driven ETV2, SOX7 and Nr2f2 and observed significant ectopic *mrc1a* expression, similar to the seven-factor pool (Fig. 3c-d and Extended Data Figs. 6e-f and 7a). Vessels ectopically expressing *mrc1a* had a sinusoidal-like morphology similar to CHT ECs and were functionally integrated into the circulatory system (Extended Data Fig. 6f and g and Supplementary Video 2). Injected embryos similarly showed ectopic expression of *sele*, *gpr182, lgmn, stab2, ifi30, ctsla* and *hexb*, as well as the *mrc1a 125bp:GFP* transgene (Extended Data Fig. 6e and h). Ectopic *mrc1a* expression was also observed when Sox18 was substituted for SOX7, or ETS1 was substituted for ETV2 (Fig. 3c and d, Extended Data Fig. 7a and c), suggesting the factors can function interchangeably. Analysis of our single cell RNA-seq data confirmed overlapping expression of multiple factors from each family within CHT ECs (Extended Data Fig. 9a). To determine whether individual factors are required for endogenous niche endothelial formation, we used previously published morpholinos (MOs). Knockdown of *etv2* has been shown to cause early vascular abnormalities that preclude its study in later niche formation^35^. Similarly, we found that depletion of both *sox7* and *sox18* together led to early vasculature abnormalities (n=67/67 animals), although individually they showed no change in *mrc1a 125bp:GFP* expression (n=90 and 104 animals, respectively). MOs targeting each of the *nr2f* family members expressed in CHT ECs (*nr2f1a*, *nr2f2* and *nr2f5*) showed no effect individually, but a low dose combination of all three led to a reduction in *mrc1a 125bp:GFP* expression and fewer HSPCs in the CHT (Extended Data Fig. 9b-c). Together these results indicate functional redundancy of the factors for both niche EC reprogramming activity and endogenous niche formation. In the mouse fetal liver (E14-17) and adult bone marrow ECs, multiple factors from the Ets, Sox and NHR families were expressed, with the highest being *Ets1, Sox18* and *Nr2f2* (Supplementary Table 7), consistent with the notion that a combination of redundant factors, with at least one from each family (although the specific factors may vary in different tissues and contexts) is a conserved feature of the vascular hematopoietic niche.

To evaluate the contribution of individual transcription factors in our 3-factor overexpression experiments we injected each factor alone. No single factor alone gave significant ectopic expression, except for ETV2, which led to ectopic expression of *mrc1a*, though at a lower frequency than with SOX7 and Nr2f2; both of which were required for optimal induction with the ETV2, SOX7 and Nr2f2 combination (Fig. 3d and Extended Data Figs. 7a-c). Some of the original seven factors, including *nr2f2* and *ets1*, had endogenous expression in the dorsal tail (Extended Data Fig. 7d), and in the majority of animals injected with ETV2 alone, ectopic expression was restricted to the dorsal tail region, suggesting the exogenous ETV2 likely works in conjunction with endogenous factors in this region. Human ETV2 alone also induced endogenous zebrafish *sox7*, *sox18*, *fli1a* and *etv2* in the dorsal tail region (Extended Data Fig. 7d). By comparison, ectopic expression with the three-factor combinations was much more widespread and in many tissues, including the anterior head and yolk regions (53% (n=338/639) of three-factor injections had ectopic yolk expression compared to 22% (n=57/265) of ETV2 alone injections). Each of the seven factors themselves had associated regions of chromatin uniquely accessible in the CHT EC fraction, harboring Ets, SoxF and NHR sites (Supplementary Table 6). Thus, overexpression of three-factor combinations likely establishes a reprogramming auto-regulatory loop that drives the niche EC program and underlies the optimal activity of the three-factor combinations to robustly induce the niche EC program.

As ectopic CHT-like ECs were frequently observed in the dorsal tail, we performed time-lapse analysis of *ubi*-driven 3-factor injected *mrc1a 125bp:GFP* and *kdrl:mCherry* embryos to determine whether these were outgrowths of CHT vasculature or derived from other cell populations in the embryo. In time-lapse movies, ectopic regions were generated independent of the CHT, often prior to the specification of the endogenous CHT ECs (Supplementary Video 1). Many of the ectopic GFP+ cells appeared to be muscle progenitors based on their size, shape and location, and these cells often underwent dramatic morphological changes – developed protrusions, became migratory and integrated into the vasculature (Supplementary Videos 3-6; Extended Data Fig. 8a). Not all cell types exhibited the same behaviors, however. Skin cells and neurons never underwent morphological changes, despite ectopically expressing the *mrc1a 125bp:GFP* transgene (Extended Data Fig. 8b). These observations are consistent with studies showing that muscle progenitors in the zebrafish embryo are susceptible to reprogramming to an endothelial fate by *etv2* overexpression^38^. As the *ubi* promoter is active very early in development, we sought to evaluate whether niche ECs could be induced at a later stage of development. We injected constructs with ETV2, SOX7 and Nr2f2 downstream of a heat shock promoter (*hsp70l*). Heat shock induction at 24 hpf resulted in large patches of niche EC gene expression throughout the animal; by 48 hours post-heat shock, these cells incorporated into the vasculature (Extended Data Fig. 8c). To test whether the CHT EC program could be induced specifically in muscle cells, we used the muscle-specific *mylz2* promoter to drive expression of the 3-factor pool. We observed *mrc1a 125bp:GFP^+^* muscle cells co-expressing the vascular marker *kdrl:mCherry*, often undergoing morphological changes (n=33/60 animals; Fig. 3e). To test whether the CHT EC program could be induced in non-CHT ECs at later stages of development, we overexpressed the transcription factors using the pan-endothelial *nrp1b* enhancer that we isolated. In these animals we observed ectopic *mrc1a 125bp:GFP* expression in non-CHT ECs, including AECs (n=22/87 animals; Fig. 3e and Extended Data Fig. 8d). The amount of ectopic expression per embryo with the *mylz2* and *nrp1b* drivers was noticeably less than with the *ubi-*driven factors, likely reflective of these drivers turning on later, in more differentiated cell types (differentiated muscle and vasculature, respectively; Fig 3e). Collectively, these data indicate that the minimal combination of Ets, SoxF and NHR factors is sufficient to induce CHT EC gene expression in multiple cell types at different stages of development, with muscle progenitors in particular being susceptible to trans-differentiation into CHT-like ECs.

### Ectopic vascular regions recruit HSPCs and support their proliferation

We next asked whether the ectopic CHT-like ECs were capable of recruiting and supporting HSPCs. Injection of 3-factor pools (ETV2 or ETS1 with SOX7 and Nr2f2) under control of the *ubi* promoter resulted in *runx1^+^* HSPCs localized outside of the CHT (Fig. 4a and b, and Extended Data Fig. 10a). 12/22 embryos with ectopic vascular patches of *mrc1a 1.3kb:GFP* had *runx1:mCherry^+^* HSPCs, often multiple, localized within the ectopic regions. In contrast, only 5/48 control embryos had HSPC localization outside the CHT. The *ubi-*induced ectopic *mrc1a:GFP^+^* ECs often formed pockets around the HSPCs and associated with *cxcl12a:DsRed2*^+^ stromal cells and *mpeg1:mCherry*+ macrophages, similar to CHT ECs (Fig. 4b, Extended Data Fig. 10c and d, and Supplementary Videos 7-9). Stromal cells or HSPCs were not localized to ectopic CHT-like vessels in the head or over the yolk (Extended Data Fig. 10e), suggesting specificity in the anatomical location and/or a functional requirement of the stromal cells in the tail for niche activity.

**Figure 4.**
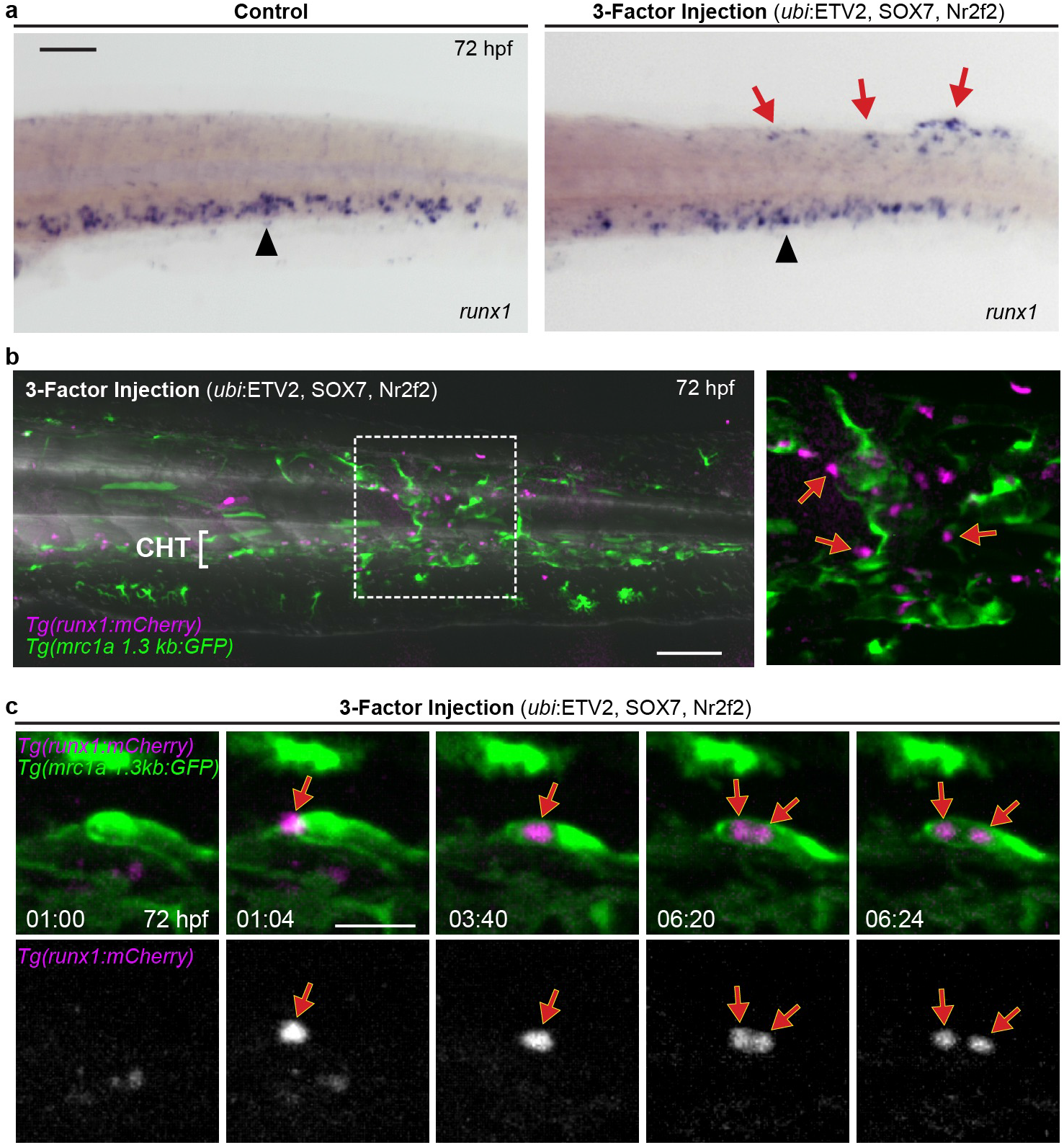
Ectopic vascular regions recruit HSPCs and support their proliferation. **a**, WISH for *runx1* shows HSPC localization in a control (left) and 3-factor injected embryo (right). Black arrowheads denote endogenous CHT localization; red arrows point to ectopic localization. **b**, Image shows *runx1:mCherry*^+^ HSPCs localized outside the CHT within a large ectopic region of *mrc1a l.3kb:GFP* expression in an embryo injected with a pool of *ubi:ETV2*, SOX7 and Nr2f2. Magnification of boxed area is shown on the right. **c**, Time-lapse series shows a *runx1:mCherry*^+^ HSPC initially arriving at an ectopic site and subsequently dividing (images show magnification of region with red arrow in Supplementary Video 8). Time is shown as hh:mm. Scale bars represent 100 μm in **a** and **b**, and 30 μm in **c**.

To investigate the dynamics of HSPC localization outside of the CHT, we used time-lapse microscopy. In control embryos the majority of HSPCs observed outside the CHT were transiently localized and only one division was observed in 10 embryos (Extended Data Fig. 10b). In 3-factor injected embryos, HSPCs localized to ectopic CHT-like ECs for several hours and often divided (6 divisions observed in 10 embryos; 5/6 divisions corresponded to HSPCs with residency times over 2.5 hours; Extended Data Fig. 10b), similar to HSPCs in the endogenous CHT^20^. We visualized recruitment, lodging and division of HSPCs, but did not observed HSPC formation at the ectopic sites (Fig. 4c and Supplementary Video 8). When HSPCs divided, daughter cells migrated away and entered circulation, presumably traveling to subsequent niches (Supplementary Video 9). Thus, multiple HSPC behaviors normally restricted to the endogenous CHT were exhibited in the ectopic patches of CHT-like ECs. Together, these data demonstrate that a minimal combination of Ets, Sox and NHR factors can induce ectopic niche ECs that associate with *cxcl12a*+ stromal cells and support the recruitment, maintenance and division of HSPCs outside the endogenous niche.

## Discussion

Our data support a model in which Ets, Sox and NHR factors specify the identity and capacity of vascular niche ECs to choreograph self-renewal and differentiation of blood stem cells. This is a conserved feature of the hematopoietic niche across species and development. The niche endothelial signature identified here includes genes that regulate adhesive interactions between ECs and circulating cells (e.g., the adhesion receptor E-selectin^25,32^ and the scavenger receptors *mrc1a, stab1* and *stab2*^31,40,41^). Another CHT niche EC gene, *gpr182*, was recently shown to be a vascular niche-expressed receptor that maintains HSPC homeostasis in fish and mice^42,43^. Our studies identified other *bona fide* niche factors including *vcam1b* and *cxcl12a* as being expressed by the niche ECs. In the bone marrow niche these genes are expressed by multiple cell types, including ECs, macrophages and mesenchymal stromal cells. This appears to be the same in the CHT niche as *cxcl12a* also marks mesenchymal stromal cells^44^ and *vcam1b* was recently reported to function in macrophages within the CHT niche^16^. There are numerous genes identified by this study that were not previously associated with the HSPC niche, including several with activities related to endocytosis and membrane trafficking: *ap1b1*, *dab2*, *pxk*, *exoc3l2a* and *snx8*. ECs in the CHT were recently shown to be highly endocytic^45^ – this activity might regulate ligand/receptor turnover or may clear potentially harmful agents, such as waste products; modified proteins; or microbial material from the niche.

Recent analyses of the mammalian HSPC niche comparing gene expression between SECs and AECs^10,46,47^ shows overlap between our niche EC signature and venous SECs in mouse bone marrow. Although AECs may also support hematopoiesis^8,9,48^, our work illustrates the capacity of SECs to recruit HSPCs and support their division. Stress-induced extramedullary hematopoiesis may involve induction of this SEC niche program. A number of our niche EC genes were enriched in adult mouse liver ECs (Fig. 1e), suggesting the liver may be ‘primed’ to support hematopoiesis under stress. Our work here highlights shared gene expression between niche ECs and head lymphatic vessels in the zebrafish embryo. Although these head lymphatic vessels in the fish do not support HSPCs, it was recently shown that lymphatic vessels are a supportive component of the hair follicle stem cell niche^49^.

Our overexpression studies indicate that 3-factor combinations of Ets, SoxF and NHR transcription factors – where specific family members are interchangeable – are able to induce niche EC gene expression in the early embryo and in more differentiated cell types, including AECs, muscle and ectodermal lineages (skin cells and neurons). Some cell populations appear to be more refractory to the developmental reprogramming, while others (e.g. muscle progenitors) trans-differentiate into CHT-like ECs with functional niche properties. Such reprogrammed niche ECs might be used in conjunction with reprogrammed stromal cells^50^ to enhance the maintenance or production of HSPCs *in vitro.* Parabiosis experiments indicate that niche size determines HSPC number^51^, and functional ectopic niches, termed ossicles, have been used to assemble a bone marrow equivalent upon transplantation^52,53^; it is likely that these structures contain SECs. These studies establish the concept that HSPC numbers could be supported *in vivo* using a reprogramming-based niche therapy to generate ectopic vascular niches at new safe harbor locations in the body, particularly for blood diseases like myelofibrosis in which the normal niche no longer functions properly. Collectively, our work advances fundamental understanding of the vascular niche that choreographs homeostasis and regeneration of blood stem cells, which may guide new strategies to culture and expand HSPCs for transplantation.

## Methods

### Animal models

Wild-type AB, *casper* or *casper*-EKK, and transgenic lines *cd41:EGFP*^54^, *runx1:mCherry* [*runx1+23:NLS-mCherry*]^17^, *kdrl(flk1):GFP* [*kdrl:GRCFP*]^55^, *kdrl:mCherry* [*kdrl:Hsa.hras-mCherry]^56^*, *cxcl12a(sdf1a):DsRed2*^44^, and *mpeg1:mCherry*^57^ were used in this study. Alternative gene names are listed in parenthesis and full transgene names are listed in brackets. All animals were housed at Boston Children’s Hospital and handled according to approved Institutional Animal Care and Use Committee (IACUC) of Boston Children’s Hospital protocols.

### Genomic analyses

For RNA tomography (tomo-seq), 72 hpf embryos were euthanized by tricaine overdose and the portion of the tail containing the CHT was manually dissected using a scalpel. The tissue was oriented in OCT tissue freezing media (Leica) in a cryomold (Tissue-TEK) with the ventral side facing the bottom of the mold. After snap freezing on dry ice, 40 individual 8 μm-thick cryosections were collected along the dorsal-ventral axis using a cryostat. The RNA from individual cryosections was extracted using TRIzol and then barcoded during a reverse transcription step prior to pooling for library preparation and sequencing according to the previously published protocol^27^. For inDrops single cell and bulk RNA-seq, *kdrl*:GFP embryos were dissociated using Liberase (Roche) and GFP^+^ cells were isolated by FACS. For bulk RNA-seq, total RNA was isolated using TRIzol LS and GenElute LPA (Sigma) carrier as per manufacturer’s instructions. Libraries were prepared from 50 ng of total RNA/sample as input using Ribogone and a SMARTer Universal Low Input RNA Kit (Clontech). For inDrops, approximately 2,000 cells were encapsulated and libraries were prepared for sequencing as previously described^58^. For ATAC-seq, embryos were dissociated using Liberase and a minimum of 12,000 cells (max 50,000) were isolated by FACS. Cells were subsequently lysed and isolated nuclei were incubated in a transposition reaction according to the published protocol^59^. All sequencing was done using an Illumina Hiseq 2500. For RNA-seq, quality control was performed by Fast QC and Cutadapt to remove adaptor sequences and low quality regions. High-quality reads were aligned to UCSC build danRer7 of the zebrafish genome using Tophat 2.0.11^60^ without novel splicing form calls. Transcript abundance and differential expression were calculated with Cufflinks 2.2.1^61^. FPKM values were used to normalize and quantify each transcript. For ATAC-seq, reads were aligned to UCSC build danRer7 of the zebrafish genome using Bowtie2 (version 2.2.1)^62^ with the following parameters: --end-to-end, -N0-, -L20. The MACS2 (version 2.1.0) peak finding algorithm^63^ was used to identify regions of ATAC-seq peaks with the following parameters: --nomodel --shift −100 --extsize 200. An initial q-value threshold of enrichment of 0.05 was used for peak calling and a more stringent q-value of 14 was used to identify peaks that were distinct between different samples. Genome-wide motif enrichment analysis was performed using HOMER^64^ and motif annotation was done using Consite^65^. Gene expression analysis of the adult kidney marrow was performed using publicly available data (https://molpath.shinyapps.io/zebrafishblood/)^66^.

### Whole mount in situ hybridization (WISH)

*In situ* hybridization was performed using a standard protocol^67^. Embryos were subsequently transferred to glycerol for scoring and imaging. *In situ* probes were generated by PCR amplification using a cDNA or plasmid (for transcription factors from other species) template followed by reverse transcription with digoxigenin-linked nucleotides. Primer sequences for all WISH probes used in this paper are provided in Supplementary Table 8.

### Transgenesis and enhancer-GFP reporter assays

Transgenic lines were established as previously described^68^. For the *mrc1a 1.3kb:GFP* and *sele 5.3kb:GFP* transgenes, 1.3 kb and 5.3 kb sequences, respectively, upstream of the transcriptional start site were PCR amplified off of genomic DNA and then TOPO-TA cloned into a p5E Gateway vector (Invitrogen), which was then recombined with GFP and a polyA tail, all flanked by Tol2 sites. For the 125 bp *mrc1a* and 158 bp *sele* enhancers, the elements were PCR amplified off of genomic DNA, TOPO-TA cloned into a p5E Gateway vector and then recombined with the mouse *Beta-globin* minimal promoter^17^ fused to GFP with a polyA tail, all flanked by Tol2 sites. Embryos were injected at the one cell-stage with Tol2 RNA and at least two independent lines showing similar expression were established for each construct: (Tg(*mrc1a 1.3kb*:GFP); Tg(*sele 5.3kb*:GFP); Tg(*mrc1a 125bp*:GFP); and Tg(*sele 158bp:GFP).* The CHT EC and pan-EC ATAC-seq elements were similarly amplified by PCR using genomic DNA and then fused to the *Beta-globin* minimal promoter and GFP. Mutational variants of 125 bp *mrc1a* and 158 bp *sele* were generated by annealing overlapping oligos followed by a T4 DNA polymerase reaction to generate blunt-ended products, which were subsequently cloned into p5E Gateway vectors (following A-tailing with Klenow Fragment (NEB)) using the same work flow as for the ATAC-seq elements. Transcription factor binding motifs were disrupted by changing nucleotides in the core binding sites, purines for pyrimidines and vice versa. Injected F0 embryos were scored between 60-72 hpf. Control and experimental groups were blinded prior to scoring and all experiments were performed at least three times, with independent clutches. GFP expression in CHT ECs or pan-EC expression was scored as significant if it was observed in at least 10% of F0 injected embryos. Embryos scored as negative had either no GFP expression or had only sparse ectopic expression in muscle cells. The sequences for primers used to amplify the *mrc1a* and *sele* regulatory elements, as well as the 15 CHT EC and 6 pan-EC ATAC-seq elements, are provided in Supplementary Table 9. The sequences for the overlapping oligos that were used to generate the enhancer variants are provided in Supplementary Table 10. The fidelity of all constructs was confirmed by sequencing prior to injection.

### Transcription factor overexpression studies

For transcription factor overexpression studies, the open reading frames for the human (FLI1, ETV2, ETS1, SOX7 and RXRA), xenopus (Sox18) or zebrafish (Nr2f2) genes were cloned into a pME Gateway vector (Invitrogen) and then recombined with the zebrafish *ubi* promoter^68^, *hsp70l* promoter^69^, *nrp1b* enhancer, or *mylz2* promoter^70^, and a polyA tail, all flanked by Tol2 sites. The fidelity of all constructs was confirmed by sequencing prior to injection. Embryos were injected with transcription factor pools (1 nl at 25 ng/μl total DNA, plus Tol2 RNA) at the one cell-stage and then screened between 24-72 hpf for ectopic niche endothelial gene expression or ectopic HSPC localization. For control and single-factor injections, the empty Tol2 Gateway destination vector was used as filler DNA in the injection mix. Expression of the transcription factors was confirmed by WISH using species-specific *in situ* probes. Ectopic expression was scored as vascular staining or vascular GFP expression outside the normal domain of gene expression. Control and experimental samples were blinded prior to scoring and all experiments were performed at least three times, with independent clutches.

### Microscopy and image analysis

Time-lapse microscopy was performed using a Yokogawa CSU-X1 spinning disk mounted on an inverted Nikon Eclipse Ti microscope equipped with dual Andor iXon EMCCD cameras and a climate controlled (maintained at 28.5C) motorized x-y stage to facilitate tiling and imaging of multiple specimens simultaneously. Screening of injected enhancer-GFP constructs and imaging of WISH embryos was performed using a Nikon SMZ18 stereomicroscope equipped with a Nikon DS-Ri2 camera. All images were acquired using NIS-Elements (Nikon) and processed using Imaris (Bitplane) or Adobe Photoshop software. HSPC dynamics were quantified using Imaris and vessel morphology was analyzed using the AngioTool software package in Fiji. Embryos were mounted for imaging as previously described^17^. Briefly, specimens were mounted in 0.8% LMP agarose with tricaine (0.16 mg/ml) in glass bottom 6-well plates and covered with E3 media containing tricaine (0.16 mg/ml).

### Flow cytometry, kidney marrow dissection, dissociation and histology

Embryos were prepared for FACS as previously described^17^. Briefly, embryos were chopped with a razor blade in cold PBS and then incubated in Liberase (Roche) for 20 minutes at 37°C before filtering the dissociated cells through a 40 μm mesh filter and transferring to 2% FBS. FACS was performed using a FACS Aria machine (BD Biosciences). Gates were set to select the brightest cells, using transgene positive and negative control samples as a guide and SYTOX Blue as a live/dead stain. Single cell RNA-seq analysis of FACS-sorted *kdrl:GFP*+ cells was used to supplement bulk RNA-seq to distinguish genes that were expressed by myeloid versus endothelial cells. At least 12,000 (50,000 max) cells were collected per sample for ATAC-seq experiments and at least 10,000 (300,000 max) cells per sample were collected for RNA-seq experiments. Kidney marrow was harvested from adult zebrafish by manual dissection and then dissociated using Liberase or fixed in 4% PFA (for histology) or dissociated by gentle pipetting (for live cell imaging). For histology the kidney marrow was embedded in paraffin prior to sectioning; alternating sections were stained with H&E or with an antibody to GFP. Mouse EC populations were sorted as Cd45^-^Pdpn^-^Cd31^+^ cells^30^.

### Morpholino injections

Morpholinos (MOs, Gene Tools) were diluted in water with phenol red and injected as previously described into 1- to 2-cell stage embryos^39,71,72^. For the Nr2f MO injections, each MO was injected at one third of the published dose. The sequences of the MOs used in this study were: *Standard Control MO*: 5’ - CCTCTTACCTCAGTTACAATTTATA - 3’; *sox7*-ATG MO: 5’- ACGCACTTATCAGAGCCGCCATGTG - 3’; *sox18*-ATG MO: 5’ –TATTCATTCCAGCAAGACCAACACG - 3’; *nr2f1a*-ATG MO: 5’-CCAGACGCTAACTACCATTGCCATA - 3’; *nr2f2*-ATG MO: 5’ – AGCCTCTCCACACTACCATTGCCAT – 3’ and *nr2f5-*ATG MO: 5’- CACTGATTTACTACCATTGCCATGC - 3’.

### Electrophoretic mobility shift assay

The Nr2f2 fragment was cloned into the pGEX2TK vector (GE Healthcare) to generate GST-tagged Nr2f2 and fidelity was verified by sequencing. The pGEX2TK-Nr2f2 protein plasmid was transformed into *E. coli* BL21 competent cells. Protein expression and purification were carried out as previously described^73^ and purified proteins were quantified against BSA. EMSAs were performed as previously described^73^. Probes were generated by annealing 100 pmol of sense and antisense oligonucleotides and 1-2 pmol of probe was used in each reaction. All primer and probe sequences are provided in Supplementary Table 11. Gel shift reactions were conducted at 4°C in 20% glycerol, 20 mM Tris (pH 8.0), 10 mM KCl, 1 mM DTT, 12.5 ng poly dI/C, 6.25 pmol of random, single-stranded oligonucleotides, BSA and the probe in the amount specified above. Samples involving the Nr2f2 protein were loaded on a 6% gel to resolve protein-DNA complexes. In reactions with cold competitors, 20x unlabeled probes were included in the reactions. Anti-NR2F2 (R&D Biosystems; cat # PP-H7147-00) was at the same amount of the Nr2f2 protein to obtain super-shifts.

### Statistical Analysis

For all graphs, error bars report mean ± s.e.m. One-way ANOVA analyses were followed by Dunnett’s (enhancer variant analyses) or Tukey’s (vessel analyses and HSPC budding) post hoc tests for multiple comparisons. Chi Square Test was used for comparing mrc1a 125bp:GFP expression data. To compare transcription factor injections, Chi Square Test and Fisher’s exact tests were used, with the Holm step-down process to correct for multiple comparisons. Unpaired two-tailed Student’s t-test or two-tailed Mann-Whitney tests were used to analyze HSPC counts and HSPC residency time, respectively. Data were analyzed using GraphPad Prism v.7.03, P < 0.05 was considered to be statistically significant. At least three independent biological replicates were performed for each experiment, with at least 15 animals from randomized, independent groups to ensure sufficient sample sized for statistical analysis. No data was excluded from any of the analyses.

## Extended Data

Extended Data contains 10 Extended Data Figures.

## Supplementary Information

Supplementary Information contains 9 Supplementary Videos and 11 Supplementary Tables.

## Acknowledgments

This work was supported by HHMI and NIH grants R01 HL04880, U54 DK110805, U01 HL100001, R24 DK092760, U01 Hl134812, P01 HL131477 and a Harvard Catalyst grant to L.I.Z. E.J.H. was an HHMI Fellow of the Helen Hay Whitney Foundation and is supported by NIH grant K01 DK111790. J.R.P. was supported by an ACS fellowship (127868-PF-15-128-01-DDC). B.K. is supported by NIH T32 HD060600. T.T.D is supported by NIH grant T32 HL098049 and an AHA Postdoctoral Fellowship. E.C.B is supported by NIH R01 AI130471 and award I01 BX-002919 from the Dept. of Veterans Affairs. We gratefully acknowledge the aquatics research staff at BCH, in particular, K. Maloney, S. Hurley and C. Lawrence. We thank R. Mathieu and M. Paktinat in the BCH flow cytometry core and M. Chatterjee in the Single Cell Core at Harvard for help with inDrops single cell RNA-seq. We thank J. Henninger for assistance with kidney marrow dissections and E. Fast and A. McConnell for comments on the manuscript.

## Author Contributions

E.J.H. designed the study, funded the project, performed experiments, managed the project, interpreted the data and wrote the manuscript. J.R.P. designed the study, performed experiments, managed the project, interpreted the data and edited the manuscript. R.J.F. S.J.W., C.M., I.F, M.L.D, C.D., T.H., M.J.F, J.K., R.R., B.L., D.A.V.E.R., K.E, E.L.H., H.G.W., S.E.R., S.H.C., B.K., J.M.G.S., T.T.D., J.P. and J.P.J performed experiments and provided technical support. A.L., S.Y., Y.Z. provided bioinformatics support. H.A.F provided biostatistics support. E.C.B, A.v.O., J.P.J and S.R. funded and supervised the project. L.I.Z. designed the study, funded and supervised the project, interpreted the data and edited the manuscript. All authors reviewed the manuscript.

## Accession Numbers

The GEO accession number for the mammalian genomic data reported in this paper is GSE100910. The zebrafish genomic data reported in this paper has been submitted to the NCBI Gene Expression Omnibus and accession numbers will be forwarded upon receipt.

## Materials & Correspondence

Correspondence and material requests should be addressed to Leonard Zon.

## References

1 Orkin, S. H. & Zon, L. I. Hematopoiesis: an evolving paradigm for stem cell biology. Cell 132, 631–644, doi:10.1016/j.cell.2008.01.025 (2008).

2 Ding, L., Saunders, T. L., Enikolopov, G. & Morrison, S. J. Endothelial and perivascular cells maintain haematopoietic stem cells. Nature 481, 457–462, doi:10.1038/nature10783 (2012).

3 Heissig, B. et al. Recruitment of stem and progenitor cells from the bone marrow niche requires MMP-9 mediated release of kit-ligand. Cell 109, 625–637 (2002).

4 Kiel, M. J. et al. SLAM family receptors distinguish hematopoietic stem and progenitor cells and reveal endothelial niches for stem cells. Cell 121, 1109–1121, doi:10.1016/j.cell.2005.05.026 (2005).

5 Perlin, J. R., Sporrij, A. & Zon, L. I. Blood on the tracks: hematopoietic stem cell-endothelial cell interactions in homing and engraftment. Journal of molecular medicine 95, 809–819, doi:10.1007/s00109-017-1559-8 (2017).

6 Sugiyama, T., Kohara, H., Noda, M. & Nagasawa, T. Maintenance of the hematopoietic stem cell pool by CXCL12-CXCR4 chemokine signaling in bone marrow stromal cell niches. Immunity 25, 977–988, doi:10.1016/j.immuni.2006.10.016 (2006).

7 Wei, Q. & Frenette, P. S. Niches for Hematopoietic Stem Cells and Their Progeny. Immunity 48, 632–648, doi:10.1016/j.immuni.2018.03.024 (2018).

8 Itkin, T. et al. Distinct bone marrow blood vessels differentially regulate haematopoiesis. Nature 532, 323–328, doi:10.1038/nature17624 (2016).

9 Kunisaki, Y. et al. Arteriolar niches maintain haematopoietic stem cell quiescence. Nature 502, 637–643, doi:10.1038/nature12612 (2013).

10 Xu, C. et al. Stem cell factor is selectively secreted by arterial endothelial cells in bone marrow. Nature communications 9, 2449, doi:10.1038/s41467-018-04726-3 (2018).

11 Avecilla, S. T. et al. Chemokine-mediated interaction of hematopoietic progenitors with the bone marrow vascular niche is required for thrombopoiesis. Nature medicine 10, 64–71, doi:10.1038/nm973 (2004).

12 Hooper, A. T. et al. Engraftment and reconstitution of hematopoiesis is dependent on VEGFR2-mediated regeneration of sinusoidal endothelial cells. Cell stem cell 4, 263–274, doi:10.1016/j.stem.2009.01.006 (2009).

13 Kim, C. H. Homeostatic and pathogenic extramedullary hematopoiesis. Journal of blood medicine 1, 13–19, doi:10.2147/JBM.S7224 (2010).

14 Murayama, E. et al. Tracing hematopoietic precursor migration to successive hematopoietic organs during zebrafish development. Immunity 25, 963–975, doi:10.1016/j.immuni.2006.10.015 (2006).

15 Wiley, D. M. et al. Distinct signalling pathways regulate sprouting angiogenesis from the dorsal aorta and the axial vein. Nature cell biology 13, 686–692, doi:10.1038/ncb2232 (2011).

16 Li, D. et al. VCAM-1(+) macrophages guide the homing of HSPCs to a vascular niche. Nature, doi:10.1038/s41586-018-0709-7 (2018).

17 Tamplin, O. J. et al. Hematopoietic stem cell arrival triggers dynamic remodeling of the perivascular niche. Cell 160, 241–252, doi:10.1016/j.cell.2014.12.032 (2015).

18 Xue, Y. et al. A 3D Atlas of Hematopoietic Stem and Progenitor Cell Expansion by Multi-dimensional RNA-Seq Analysis. Cell reports 27, 1567–1578 e1565, doi:10.1016/j.celrep.2019.04.030 (2019).

19 Murayama, E. et al. NACA deficiency reveals the crucial role of somite-derived stromal cells in haematopoietic niche formation. Nature communications 6, 8375, doi:10.1038/ncomms9375 (2015).

20 Blaser, B. W. et al. CXCR1 remodels the vascular niche to promote hematopoietic stem and progenitor cell engraftment. The Journal of experimental medicine 214, 1011–1027, doi:10.1084/jem.20161616 (2017).

21 Butler, J. M. et al. Endothelial cells are essential for the self-renewal and repopulation of Notch-dependent hematopoietic stem cells. Cell stem cell 6, 251–264, doi:10.1016/j.stem.2010.02.001 (2010).

22 Ding, L. & Morrison, S. J. Haematopoietic stem cells and early lymphoid progenitors occupy distinct bone marrow niches. Nature 495, 231–235, doi:10.1038/nature11885 (2013).

23 Greenbaum, A. et al. CXCL12 in early mesenchymal progenitors is required for haematopoietic stem-cell maintenance. Nature 495, 227–230, doi:10.1038/nature11926 (2013).

24 Mahony, C. B., Fish, R. J., Pasche, C. & Bertrand, J. Y. tfec controls the hematopoietic stem cell vascular niche during zebrafish embryogenesis. Blood 128, 1336–1345, doi:10.1182/blood-2016-04-710137 (2016).

25 Winkler, I. G. et al. Vascular niche E-selectin regulates hematopoietic stem cell dormancy, self renewal and chemoresistance. Nature medicine 18, 1651–1657, doi:10.1038/nm.2969 (2012).

26 Xue, Y. et al. The Vascular Niche Regulates Hematopoietic Stem and Progenitor Cell Lodgment and Expansion via klf6a-ccl25b. Developmental cell 42, 349–362 e344, doi:10.1016/j.devcel.2017.07.012 (2017).

27 Junker, J. P. et al. Genome-wide RNA Tomography in the zebrafish embryo. Cell 159, 662–675, doi:10.1016/j.cell.2014.09.038 (2014).

28 Theodore, L. N. et al. Distinct Roles for Matrix Metalloproteinases 2 and 9 in Embryonic Hematopoietic Stem Cell Emergence, Migration, and Niche Colonization. Stem cell reports 8, 1226–1241, doi:10.1016/j.stemcr.2017.03.016 (2017).

29 Zapata, A. Ultrastructural study of the teleost fish kidney. Developmental and comparative immunology 3, 55–65, doi:10.1016/s0145-305x(79)80006-3 (1979).

30 Barry, D. M. et al. Molecular determinants of nephron vascular specialization in the kidney. Nature communications 10, 5705, doi:10.1038/s41467-019-12872-5 (2019).

31 Irjala, H. et al. Mannose receptor is a novel ligand for L-selectin and mediates lymphocyte binding to lymphatic endothelium. The Journal of experimental medicine 194, 1033–1042, doi:10.1084/jem.194.8.1033 (2001).

32 Frenette, P. S., Mayadas, T. N., Rayburn, H., Hynes, R. O. & Wagner, D. D. Susceptibility to infection and altered hematopoiesis in mice deficient in both P- and E-selectins. Cell 84, 563–574 (1996).

33 Aranguren, X. L. et al. Transcription factor COUP-TFII is indispensable for venous and lymphatic development in zebrafish and Xenopus laevis. Biochemical and biophysical research communications 410, 121–126, doi:10.1016/j.bbrc.2011.05.117 (2011).

34 De Val, S. et al. Combinatorial regulation of endothelial gene expression by ets and forkhead transcription factors. Cell 135, 1053–1064, doi:10.1016/j.cell.2008.10.049 (2008).

35 Sumanas, S. & Lin, S. Ets1-related protein is a key regulator of vasculogenesis in zebrafish. PLoS biology 4, e10, doi:10.1371/journal.pbio.0040010 (2006).

36 Ginsberg, M. et al. Efficient direct reprogramming of mature amniotic cells into endothelial cells by ETS factors and TGFbeta suppression. Cell 151, 559–575, doi:10.1016/j.cell.2012.09.032 (2012).

37 Morita, R. et al. ETS transcription factor ETV2 directly converts human fibroblasts into functional endothelial cells. Proceedings of the National Academy of Sciences of the United States of America 112, 160–165, doi:10.1073/pnas.1413234112 (2015).

38 Veldman, M. B. et al. Transdifferentiation of fast skeletal muscle into functional endothelium in vivo by transcription factor Etv2. PLoS biology 11, e1001590, doi:10.1371/journal.pbio.1001590 (2013).

39 Swift, M. R. et al. SoxF factors and Notch regulate nr2f2 gene expression during venous differentiation in zebrafish. Developmental biology 390, 116–125, doi:10.1016/j.ydbio.2014.03.018 (2014).

40 Irjala, H. et al. The same endothelial receptor controls lymphocyte traffic both in vascular and lymphatic vessels. European journal of immunology 33, 815–824, doi:10.1002/eji.200323859 (2003).

41 Jung, M. Y., Park, S. Y. & Kim, I. S. Stabilin-2 is involved in lymphocyte adhesion to the hepatic sinusoidal endothelium via the interaction with alphaMbeta2 integrin. Journal of leukocyte biology 82, 1156–1165, doi:10.1189/jlb.0107052 (2007).

42 Xia, J. et al. A single-cell resolution developmental atlas of hematopoietic stem and progenitor cell expansion in zebrafish. Proceedings of the National Academy of Sciences 118, e2015748118, doi:10.1073/pnas.2015748118 (2021).

43 Le Mercier, A. et al. GPR182 is an endothelium-specific atypical chemokine receptor that maintains hematopoietic stem cell homeostasis. Proceedings of the National Academy of Sciences 118, e2021596118, doi:10.1073/pnas.2021596118 (2021).

44 Glass, T. J. et al. Stromal cell-derived factor-1 and hematopoietic cell homing in an adult zebrafish model of hematopoietic cell transplantation. Blood 118, 766–774, doi:10.1182/blood-2011-01-328476 (2011).

45 Campbell, F. et al. Directing Nanoparticle Biodistribution through Evasion and Exploitation of Stab2-Dependent Nanoparticle Uptake. ACS nano 12, 2138–2150, doi:10.1021/acsnano.7b06995 (2018).

46 Baryawno, N. et al. A Cellular Taxonomy of the Bone Marrow Stroma in Homeostasis and Leukemia. Cell 177, 1915–1932 e1916, doi:10.1016/j.cell.2019.04.040 (2019).

47 Tikhonova, A. N. et al. The bone marrow microenvironment at single-cell resolution. Nature 569, 222–228, doi:10.1038/s41586-019-1104-8 (2019).

48 Khan, J. A. et al. Fetal liver hematopoietic stem cell niches associate with portal vessels. Science 351, 176–180, doi:10.1126/science.aad0084 (2016).

49 Gur-Cohen, S. e. a. Stem cell-driven lymphatic remodeling coordinates tissue regeneration. Science 366, 1218–1225 (2019).

50 Nakahara, F. et al. Engineering a haematopoietic stem cell niche by revitalizing mesenchymal stromal cells. Nature cell biology 21, 560–567, doi:10.1038/s41556-019-0308-3 (2019).

51 Chen, J. et al. Mobilization as a preparative regimen for hematopoietic stem cell transplantation. Blood 107, 3764–3771, doi:10.1182/blood-2005-09-3593 (2006).

52 Chan, C. K. et al. Endochondral ossification is required for haematopoietic stem-cell niche formation. Nature 457, 490–494, doi:10.1038/nature07547 (2009).

53 Sacchetti, B. et al. Self-renewing osteoprogenitors in bone marrow sinusoids can organize a hematopoietic microenvironment. Cell 131, 324–336, doi:10.1016/j.cell.2007.08.025 (2007).

54 Lin, H. F. et al. Analysis of thrombocyte development in CD41-GFP transgenic zebrafish. Blood 106, 3803–3810, doi:10.1182/blood-2005-01-0179 (2005).

55 Cross, L. M., Cook, M. A., Lin, S., Chen, J. N. & Rubinstein, A. L. Rapid analysis of angiogenesis drugs in a live fluorescent zebrafish assay. Arteriosclerosis, thrombosis, and vascular biology 23, 911–912, doi:10.1161/01.ATV.0000068685.72914.7E (2003).

56 Chi, N. C. et al. Foxn4 directly regulates tbx2b expression and atrioventricular canal formation. Genes & development 22, 734–739, doi:10.1101/gad.1629408 (2008).

57 Ellett, F., Pase, L., Hayman, J. W., Andrianopoulos, A. & Lieschke, G. J. mpeg1 promoter transgenes direct macrophage-lineage expression in zebrafish. Blood 117, e49–56, doi:10.1182/blood-2010-10-314120 (2011).

58 Zilionis, R. et al. Single-cell barcoding and sequencing using droplet microfluidics. Nature protocols 12, 44–73, doi:10.1038/nprot.2016.154 (2017).

59 Buenrostro, J. D., Giresi, P. G., Zaba, L. C., Chang, H. Y. & Greenleaf, W. J. Transposition of native chromatin for fast and sensitive epigenomic profiling of open chromatin, DNA-binding proteins and nucleosome position. Nature methods 10, 1213–1218, doi:10.1038/nmeth.2688 (2013).

60 Trapnell, C., Pachter, L. & Salzberg, S. L. TopHat: discovering splice junctions with RNA-Seq. Bioinformatics 25, 1105–1111, doi:10.1093/bioinformatics/btp120 (2009).

61 Trapnell, C. et al. Transcript assembly and quantification by RNA-Seq reveals unannotated transcripts and isoform switching during cell differentiation. Nature biotechnology 28, 511–515, doi:10.1038/nbt.1621 (2010).

62 Langmead, B. & Salzberg, S. L. Fast gapped-read alignment with Bowtie 2. Nature methods 9, 357–359, doi:10.1038/nmeth.1923 (2012).

63 Zhang, Y. et al. Model-based analysis of ChIP-Seq (MACS). Genome biology 9, R137, doi:10.1186/gb-2008-9-9-r137 (2008).

64 Heinz, S. et al. Simple combinations of lineage-determining transcription factors prime cis-regulatory elements required for macrophage and B cell identities. Molecular cell 38, 576–589, doi:10.1016/j.molcel.2010.05.004 (2010).

65 Sandelin, A., Wasserman, W. W. & Lenhard, B. ConSite: web-based prediction of regulatory elements using cross-species comparison. Nucleic acids research 32, W249–252, doi:10.1093/nar/gkh372 (2004).

66 Tang, Q. et al. Dissecting hematopoietic and renal cell heterogeneity in adult zebrafish at single-cell resolution using RNA sequencing. The Journal of experimental medicine 214, 2875–2887, doi:10.1084/jem.20170976 (2017).

67 Thisse, C. & Thisse, B. High-resolution in situ hybridization to whole-mount zebrafish embryos. Nature protocols 3, 59–69, doi:10.1038/nprot.2007.514 (2008).

68 Mosimann, C. et al. Ubiquitous transgene expression and Cre-based recombination driven by the ubiquitin promoter in zebrafish. Development 138, 169–177, doi:10.1242/dev.059345 (2011).

69 Adam, A., Bartfai, R., Lele, Z., Krone, P. H. & Orban, L. Heat-inducible expression of a reporter gene detected by transient assay in zebrafish. Experimental cell research 256, 282–290, doi:10.1006/excr.2000.4805 (2000).

70 Ju, B. et al. Recapitulation of fast skeletal muscle development in zebrafish by transgenic expression of GFP under the mylz2 promoter. Developmental dynamics: an official publication of the American Association of Anatomists 227, 14–26, doi:10.1002/dvdy.10273 (2003).

71 Chen, J., Carney, S. A., Peterson, R. E. & Heideman, W. Comparative genomics identifies genes mediating cardiotoxicity in the embryonic zebrafish heart. Physiological Genomics 33, 148–158, doi:10.1152/physiolgenomics.00214.2007 (2008).

72 Hwang, S.-P. L. et al. Nuclear Receptor Subfamily 2 Group F Member 1a (nr2f1a) Is Required for Vascular Development in Zebrafish. PLoS ONE 9, e105939, doi:10.1371/journal.pone.0105939 (2014).

73 Dinh, T. T. et al. The floral homeotic protein APETALA2 recognizes and acts through an AT-rich sequence element. Development 139, 1978–1986, doi:10.1242/dev.077073 (2012).

