## Supplementary Information for "Transcription factor induction of vascular blood stem cell niches *in vivo*"

Contains 10 Extended Data Figures, 9 Supplementary Videos and 11 Supplementary Tables.

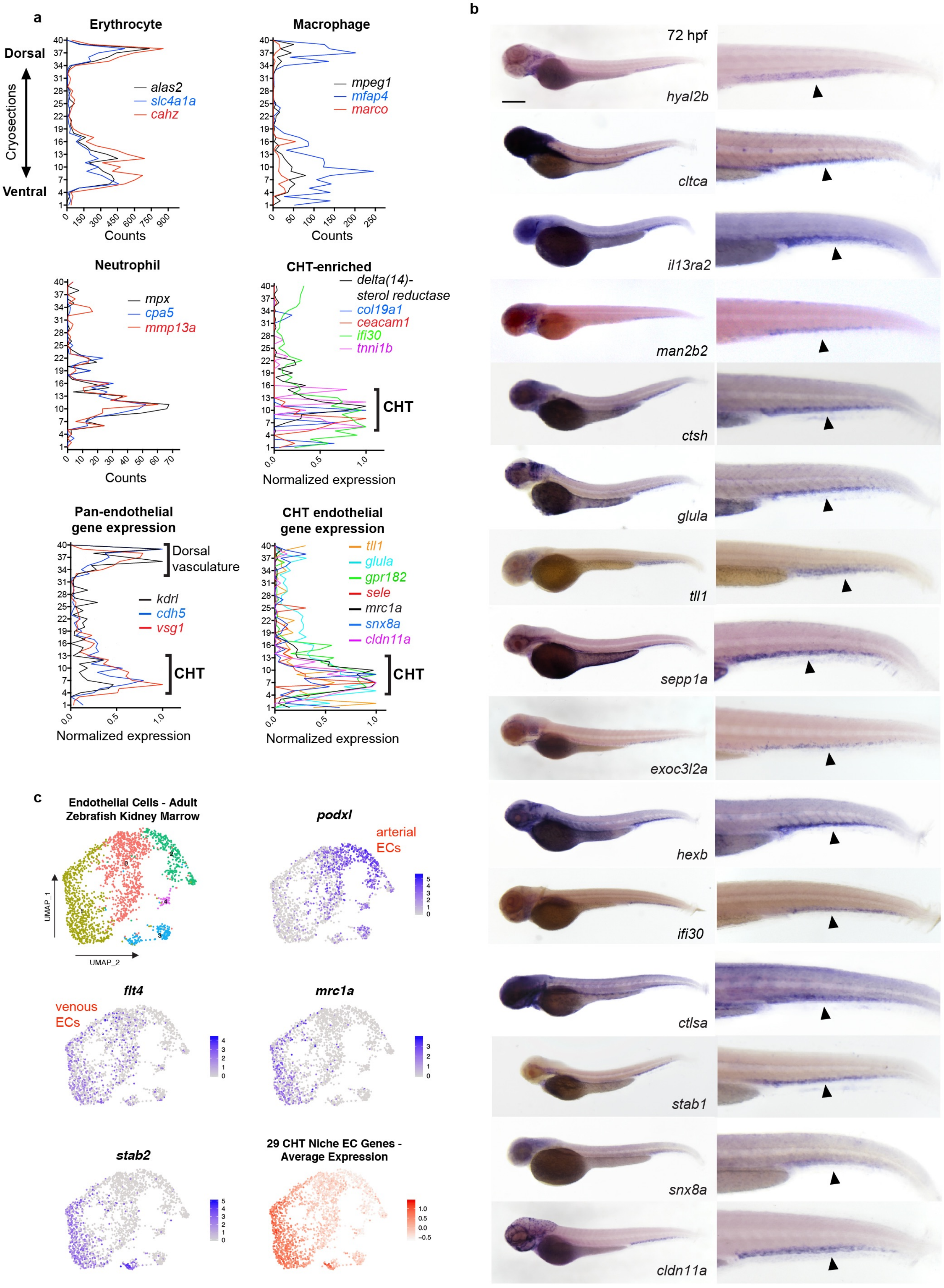


### Extended Data Figure 1

**Extended Data Figure 1 | RNA tomography and niche-specific endothelial gene expression**. **a**, Graphs show tomo-seq expression traces for individual tissue-specific genes. **b**, WISH validates the CHT-enriched expression (arrowheads) of CHT EC genes identified using a combination of tomo-seq and tissue-specific RNA-seq. Scale bars represent 250 μm in this and all subsequent Extended Data Figures unless noted otherwise. **c**, Uniform Manifold Approximation and Projection (UMAP) plots show cell clustering and gene expression from a single cell RNA-seq analysis of *kdrl:mCherry*$^{+}$ endothelial cells isolated from adult zebrafish kidney marrow. Marker genes are shown for arterial (*podxl*) and venous (*flt4*) endothelial cells, as well as CHT ECs (*mrc1a* and *stab2*). The average expression of the 29 CHT niche EC genes is shown in the bottom right plot. Spectral scales report z-scores.

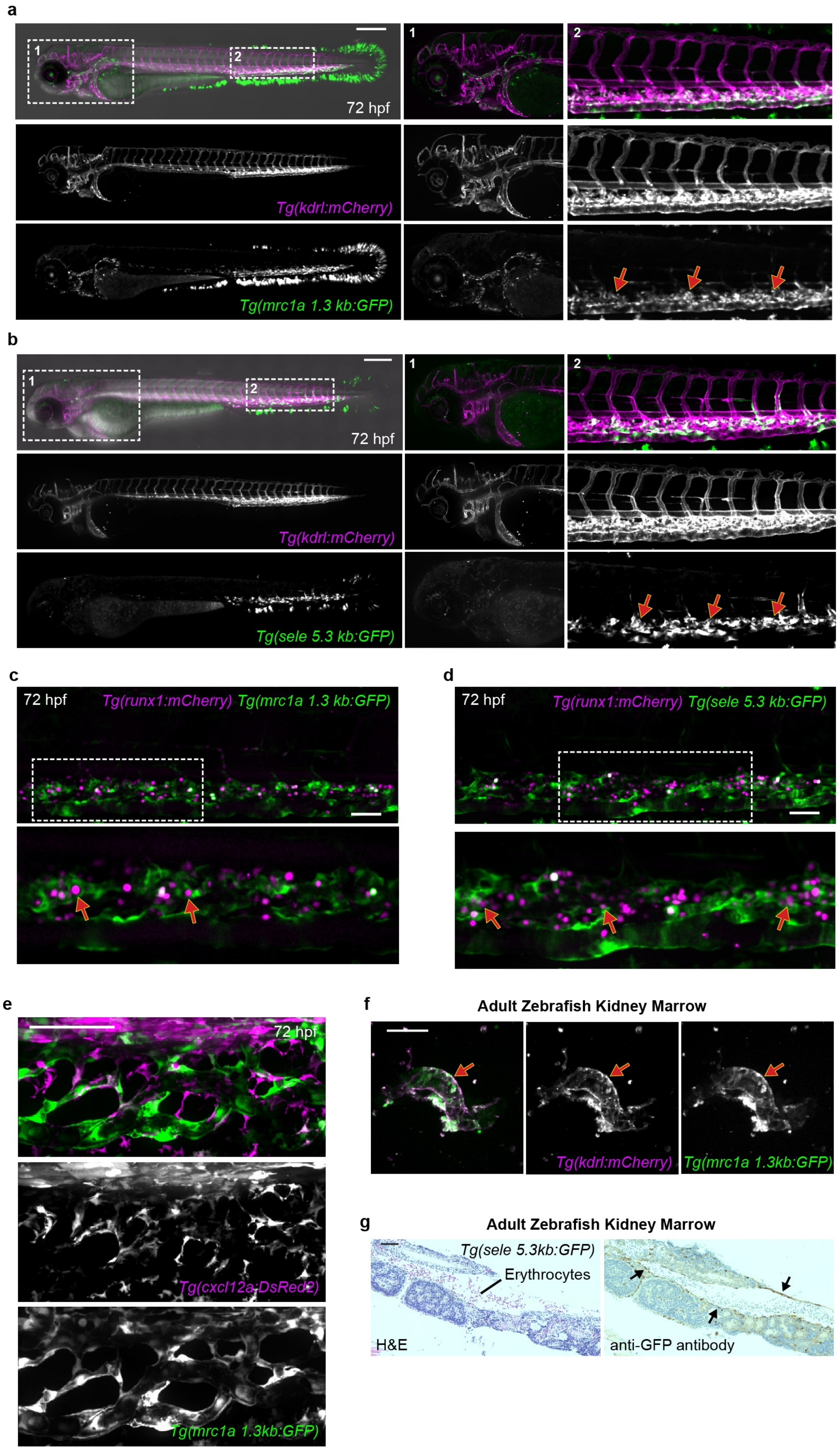


**Extended Data Figure 2**

**Extended Data Figure 2 | GFP reporter transgenes selectively label ECs in the HSPC niche**. **a**, Images show a double transgenic embryo carrying the pan-endothelial marker *kdrl:mCherry* (magenta) and the *mrc1a 1.3kb:GFP* transgene (green). Magnifications of boxed areas are shown on the right. The highest levels of vascular GFP expression are observed in CHT ECs (red arrows); while lower levels of expression are observed in the anterior head region, although some of these cells do not express the *kdrl:mCherry* transgene. **b**, Images show a double transgenic embryo carrying the pan-endothelial marker *kdrl:mCherry* (magenta) and the *sele 5.3kb:GFP* transgene (green). Magnifications of boxed areas are shown on the right. The highest levels of vascular GFP expression are observed in CHT ECs (red arrows); while lower levels of expression are observed in the anterior head region, although some of these cells do not express the *kdrl:mCherry* transgene. **c**, Images show *runx1:mCherry*$^{+}$ HSPCs (magenta) directly interacting with *mrc1a 1.3kb:GFP*$^{+}$ ECs within the CHT niche (red arrows). Bottom panel shows magnification of boxed area. **d**, Images show *runx1:mCherry*$^{+}$ HSPCs (magenta) directly interacting with *sele 5.3kb:GFP*$^{+}$ ECs within the CHT niche (red arrows). Bottom panel shows magnification of boxed area. **e**, *cxcl12a*:DsRed2$^{+}$ stromal cells (magenta) are closely associated with *mrc1a 1.3kb:GFP*$^{+}$ ECs in the CHT. **f**, Images show a segment of vasculature (red arrows) dissected from the kidney of a *mrc1a 1.3kb:GFP; kdrl:mCherry* double transgenic adult zebrafish. **g**, Images show sequential sections through an adult kidney isolated from a *sele 5.3kb:GFP* transgenic fish. Sections were stained with H&E (left) and with an antibody against GFP (right). Black arrows point to GFP$^{+}$ vascular endothelial cells. Scale bars represent 250 μm in **a**-**b**, and 50 μm in **c**-**g**.

a

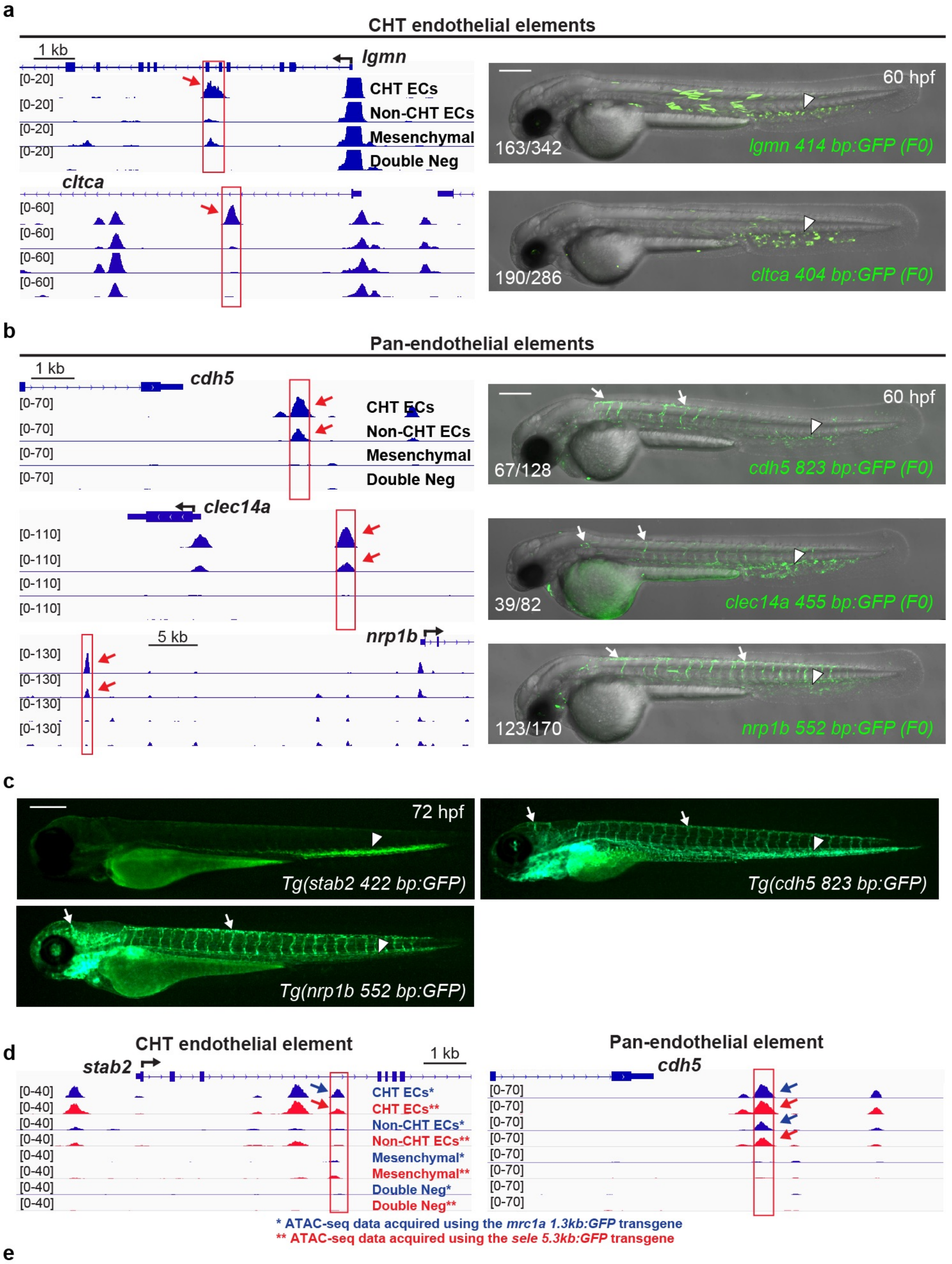


e

Motifs over-represented in CHT endothelial elements

| Rank | Motif | P-value | % of Targets | TF Family | Best Match/Details |
|---|---|---|---|---|---|
| 1 | 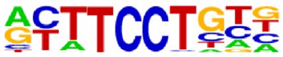 | 1e-1487 | 58.15% | Ets | ERG(ETS)/VCaP-ERG-ChIP-Seq(GSE14097)/Homer(0.948) |
| 2 | 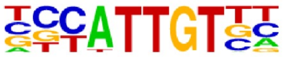 | 1e-206 | 16.95% | Sox | MA0078.1_Sox17/Jaspar(0.950) |
| 3 | 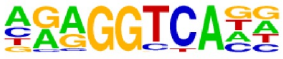 | 1e-153 | 23.79% | NHR | MF0004.1 Nuclear Receptor class/Jaspar(0.943) |

Motifs over-represented in pan-endothelial elements

| Rank | Motif | P-value | % of Targets | TF Family | Best Match/Details |
|---|---|---|---|---|---|
| 1 | 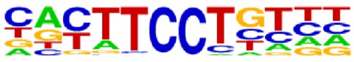 | 1e-886 | 53.67% | Ets | MA0474.1_Erg/Jaspar(0.951) |
| 2 | 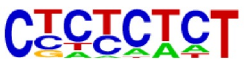 | 1e-89 | 32.09% | Zinc Finger | PRDM1(Zf)/Hela-PRDM1-ChIP-Seq(GSE31477)/Homer(0.678) |
| 3 | 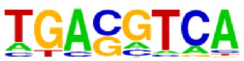 | 1e-81 | 19.22% | Leucine Zipper | Atf7(bZIP)/3T3L1-Atf7-ChIP-Seq(GSE56872)/Homer(0.940) |

### Extended Data Figure 3

**Extended Data Figure 3 | A *cis*-regulatory landscape for HSPC niche- and pan-endothelial gene expression**. **a**, Gene tracks show regions of chromatin that were uniquely open in the mCherry$^+$; GFP$^+$ CHT EC fraction (red boxes and arrows). Images on the right show embryos injected with a CHT EC enhancer-GFP reporter construct corresponding to the red boxed regions. Arrowhead points to GFP expression in CHT ECs. **b**, Gene tracks show regions of chromatin that were open in both the mCherry$^+$GFP$^+$ (CHT EC) and mCherry$^+$GFP$^-$ (non-CHT EC) populations (red boxes and arrows). Images on the right show embryos injected with pan-endothelial enhancer-GFP reporter constructs corresponding to the red boxed regions. Arrows point to GFP expression in non-CHT ECs and arrowheads point to expression in CHT ECs. **c**, Images show reporter expression of stable enhancer-GFP transgenes. Arrows point to GFP expression in non-CHT ECs and arrowheads point to expression in CHT ECs. **d**, Gene tracks show regions of chromatin that were uniquely open in the mCherry$^+$; GFP$^+$ CHT EC fraction (left; red box and arrows) or a region of chromatin open in both the mCherry$^+$GFP$^+$ (CHT EC) and mCherry$^+$GFP$^-$ (non-CHT EC) populations (right; red box and arrows). The blue tracks show ATAC-seq data obtained using the *mrc1a 1.3kb:GFP* transgene, while the red tracks show ATAC-seq data obtained using the *sele 5.3kb:GFP* transgene. **e**, Tables show the transcription factor binding motifs most enriched in CHT EC regions (top) or pan-endothelial regions (bottom).

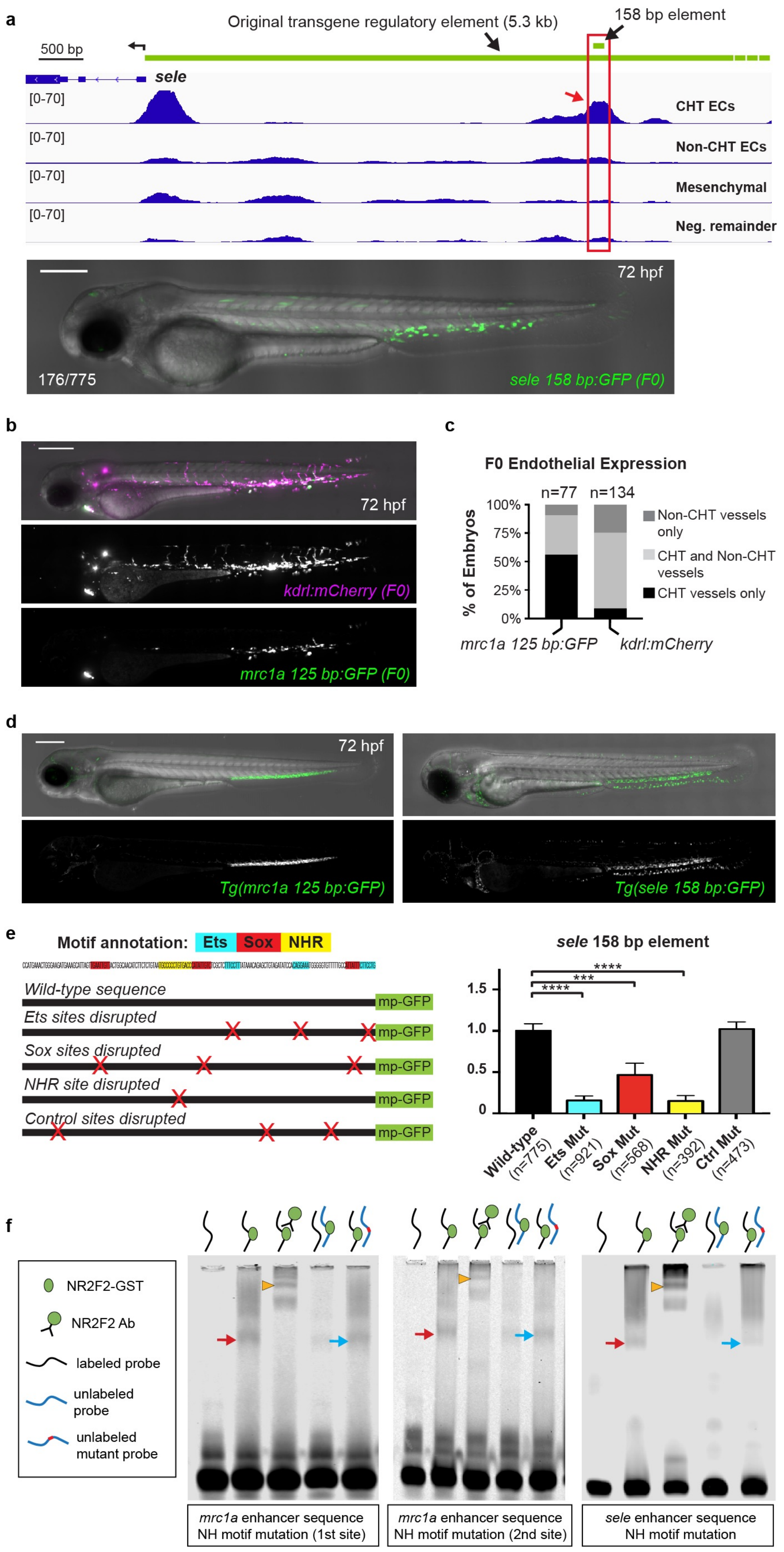


**Extended Data Figure 4**

**Extended Data Figure 4 | Enhancer sequencing for tissue-specific gene expression in niche ECs**. **a**, Gene tracks show a region of chromatin upstream of *sele* that was uniquely open in the double positive CHT EC fraction but not the other three cell populations (red box and arrow). Green bars denote the position of the 158 bp enhancer sequence and the 5.3 kb sequence used to generate the *sele:GFP* reporter transgenes. Image (bottom) shows an F0 embryo injected with *sele 158 bp:GFP.* **b**, Images show an F0 embryo injected with *mrc1a 125 bp:GFP* and *kdrl:mCherry* plasmids. mCherry expression is observed in ECs throughout the embryo, whereas GFP expression is mostly restricted to CHT ECs. **c**, Graph reports the anatomical location of endothelial expression in F0 embryos that were injected with *mrc1a 125 bp:GFP* and *kdrl:mCherry* plasmids. **d**, Images show embryos expressing the stably integrated *mrc1a 125 bp:GFP* (left) and *sele 158bp:GFP* (right) transgenes. **e**, Wild-type sequence of the 158 bp *sele* enhancer is shown, annotated with colors highlighting the Ets, Sox and NHR binding motifs (top). Schematic depicts sequence variants in which each class of motif or control regions were targeted by mutation. Red X's denote the location of targeted sites. mp-GFP: mouse *Beta-globin* minimal promoter fused to GFP. Graph reports the frequency of embryos with GFP expression in CHT ECs after injection with wild-type sequences or mutated variants of the *sele* 158 bp enhancer. Data is normalized to the wild-type control (23% $GFP^+$ CHT ECs (176/775)). Mean +/- s.e.m., One-way ANOVA with Dunnett's multiple comparisons test; ***P<0.001, ****P<0.0001. **f**, Images show electophoretic mobility shift assays with recombinant Nr2f2-GST that was incubated with DNA sequences spanning the NHR motifs present in the 125 bp *mrc1a* (left two gels) or 158 bp *sele* (right gel) enhancer sequences. Red arrows point to DNA:protein binding while orange arrowheads point to super-shifted DNA:protein complexes. Labeled DNA:protein complexes were outcompeted by unlabeled wild-type probe (lane 4) but not by unlabeled probe in which the NHR motif was disrupted by mutation (blue arrows).

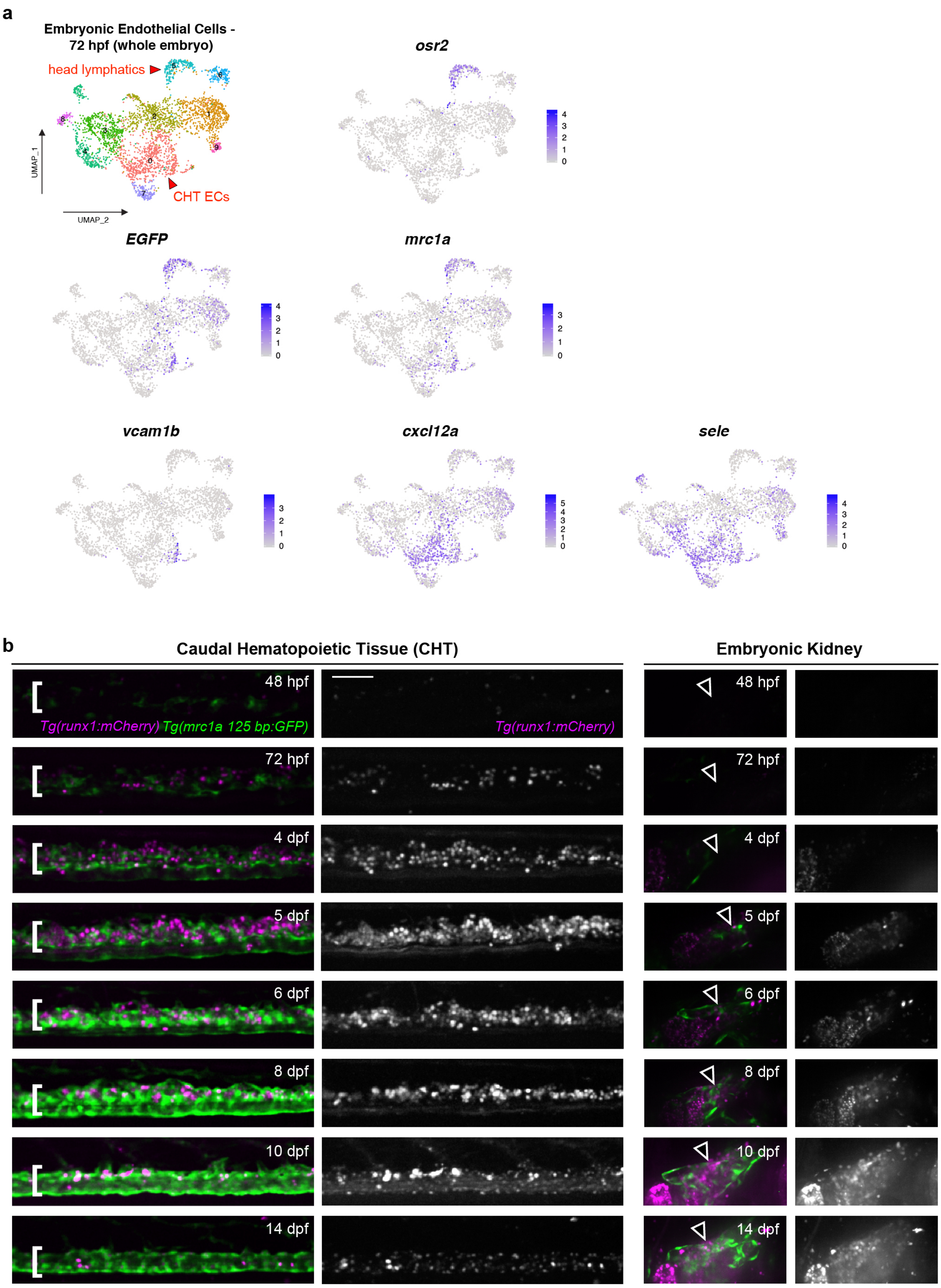


**Extended Data Figure 5**

**Extended Data Figure 5 | The *mrc1a 125bp:GFP* enhancer transgene selectively labels niche ECs**. **a**, Uniform Manifold Approximation and Projection (UMAP) plots show cell clustering and gene expression from a single cell RNA-seq analysis of endothelial cells (whole embryo) isolated from *mrc1a 125bp:GFP*; *kdrl:mCherry* double positive embryos at 72 hpf. *osr2* expression is shown as a marker of the head lymphatic EC population. Spectral scales report z-scores. **b**, Images show *mrc1a 125bp:GFP* (green) and *runx1:mCherry* (magenta) expression in the CHT (left) and kidney (right) at 8 different developmental time points. White brackets denote the location of the CHT and arrowheads point to the location of the developing kidney. Grayscale images of the *runx1:mCherry* signal are shown to the right of color overlays. Identical settings were used for image acquisition at each time point. The fluorescence intensity for the images in the figure was adjusted relative to the highest expression at 8 dpf. This results in the reduced fluorescence intensity in the image for the 48 hpf time point. Scale bar represents 100µm.

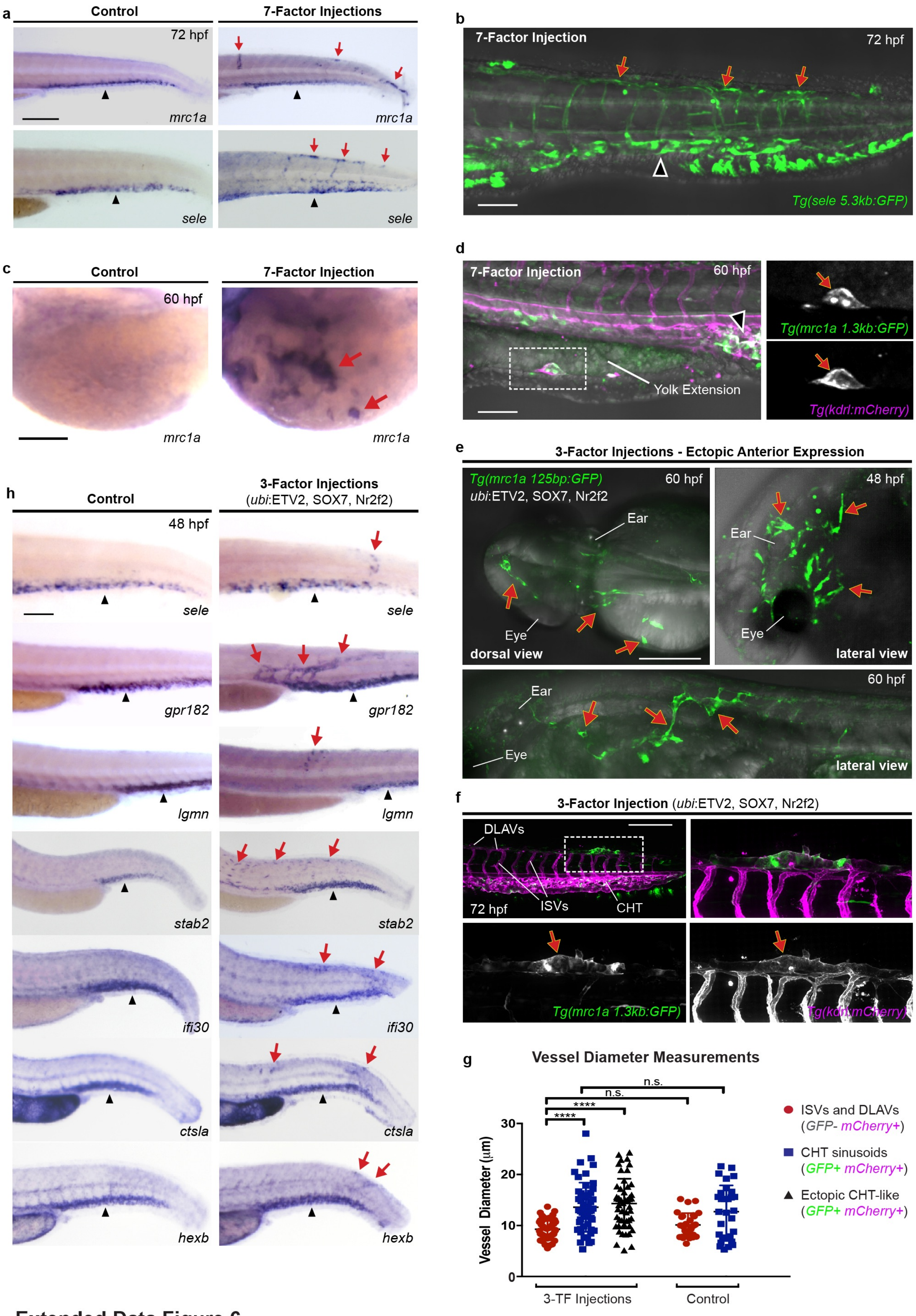


### Extended Data Figure 6

**Extended Data Figure 6 | Transcription factor overexpression induces ectopic expression of the CHT niche endothelial program**. **a**, Images show embryos that were injected with control DNA (left) or a pool of seven transcription factors (right) from the Ets, Sox and NHR families (FLI1, ETV2, ETS1, SOX7, Sox18, Nr2f2 and RXRA) and then stained by WISH for *mrc1a* (top) or *sele* (bottom). Red arrows denote regions of ectopic expression and black arrowheads point to normal domains of expression in all panels of this figure. **b**, Image shows a *sele 5.3kb:GFP* transgenic embryo that was injected with the 7-factor pool. **c**, Images show WISH staining for *mrc1a* over the yolk ball in a control (left) and 7-factor injected embryo (right). **d**, Images show ectopic expression of the *mrc1a 1.3kb:GFP* and *kdrl:mCherry* transgenes over the yolk extension in a 7-factor injected embryo. Magnification of the boxed area is shown at the right. **e**, Images show ectopic GFP expression in vessels in the anterior head and yolk region of *mrc1a 125bp:GFP* transgenic embryos injected with the 3-factor *ubi*:ETV2, SOX7, Nr2f2 mixture. **f**, Images show a large vessel in the dorsal tail region (red arrow) ectopically expressing *mrc1a 125bp:GFP* (green) and *kdrl:mCherry* (magenta). Magnification of white dotted box is shown. Dorsal longitudinal anastomotic vessels (DLAVs); Intersegmental vessels (ISVs). **g**, Graph reports quantitative measurements of vessel diameter for intersegmental vessels (ISVs) and dorsal longitudinal anastomotic vessels (DLAVs), and CHT sinusoidal vessels in *ubi*:ETV2, SOX7 and N2f2 and control injected embryos. One-way ANOVA with Tukey's test for multiple comparisons; ****$P<0.0001$; n.s. = not significant. **h**, Images show WISH for CHT niche EC genes in control embryos (left) and embryos injected with the 3-factor *ubi*:ETV2, SOX7 and

Nr2f2 combination (right). Scale bars represent 250 μm in **a**, **e**, and **f**, and 100 μm in **b-d** and **h**.

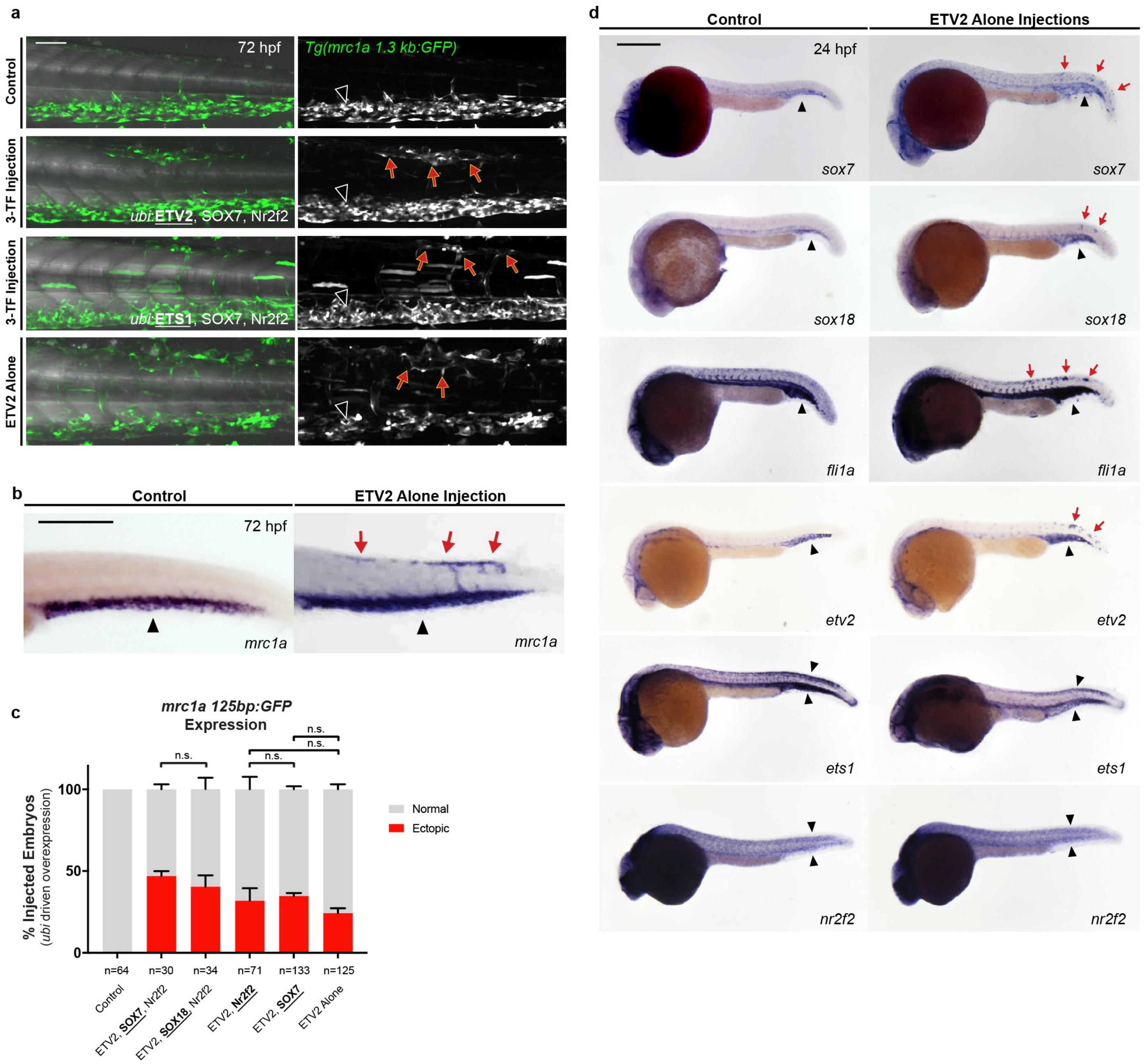


**Extended Data Figure 7**

**Extended Data Figure 7 | Transcription factor induction of niche EC gene expression**. **a**, Images show *mrc1a 1.3kb:GFP* transgenic embryos that were injected with different combinations of transcription factors at the one-cell stage. Grayscale images of the GFP signal are shown on the right. Red arrows denote regions of ectopic expression and black arrowheads point to normal domains of expression in all panels of this figure. **b**, Injection of human ETV2 alone induces ectopic expression of the endogenous *mrc1a* gene. **c**, Graph reports the percentage of transcription factor-injected embryos that showed ectopic expression of *mrc1a 125bp:GFP.* Fisher's exact test for pairwise comparisons was used; n.s. = not significant. **d**, Injection of human ETV2 alone induces ectopic expression of zebrafish transcription factors, including *sox7*, *sox18*, *fli1a* and *etv2*.

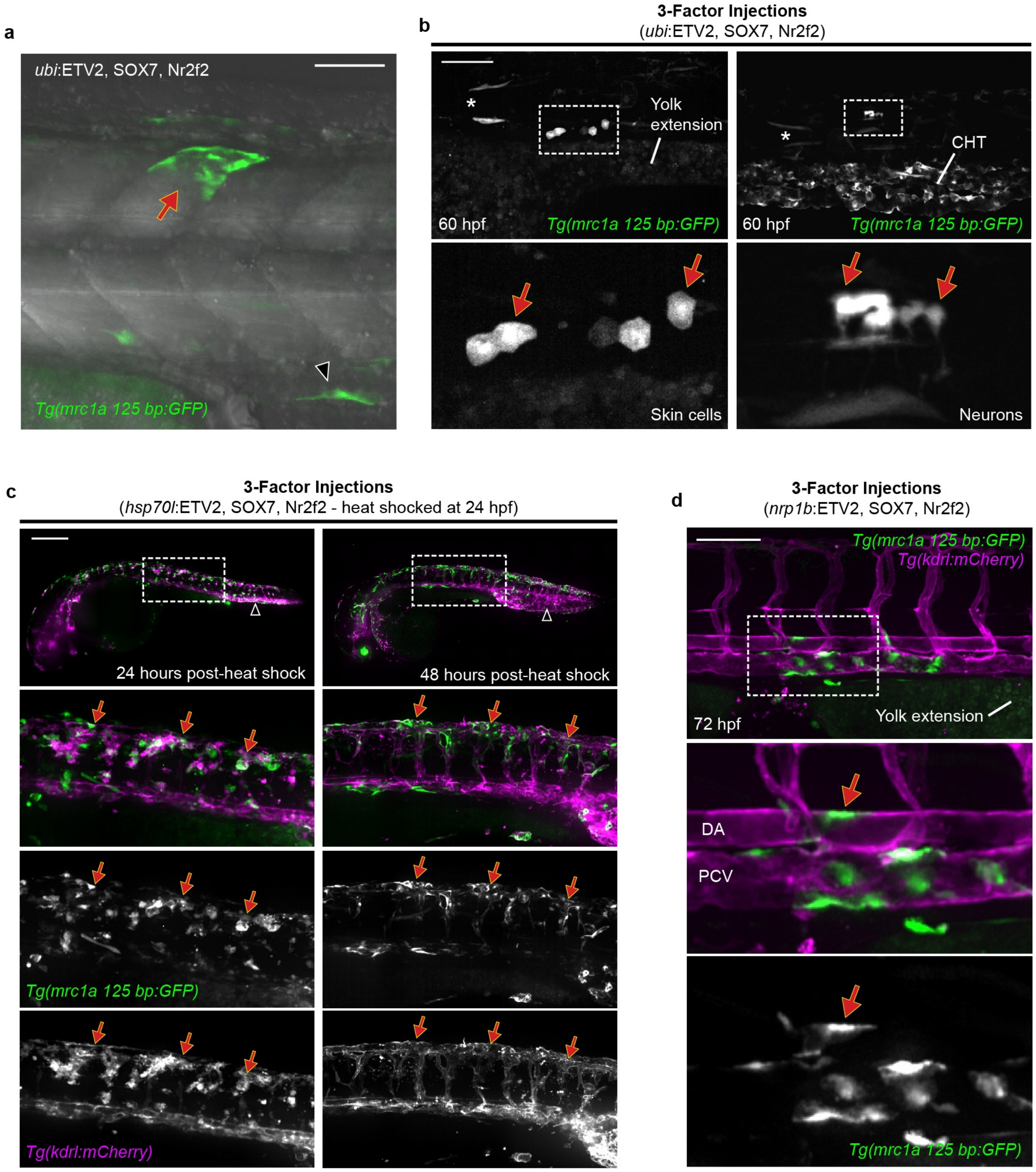


**Extended Data Figure 8**

**Extended Data Figure 8 | Tissue-specific induction of niche EC gene expression**. **a**, Images show a cluster of GFP$^+$ muscle-shaped cells ectopically expressing the *mrc1a 125 bp:GFP* transgene in a *ubi*:ETV2, SOX7, Nr2f2 injected embryo. Image corresponds to Supplementary Video 4. **b**, Images show GFP$^+$ skin cells (left) and neurons (right) ectopically expressing the *mrc1a 125 bp:GFP* transgene in a *ubi*:ETV2, SOX7, Nr2f2 injected embryo. Asterisks denote expression in muscle cells. **c**, Images show ectopic expression in *mrc1a 125bp:GFP; kdrl:mCherry* double positive embryos that were injected with *hsp70l:*ETV2, SOX7, Nr2f2 plasmids at the one-cell stage and then heat shocked at 24 hpf. Magnification of boxed regions is shown at bottom. **d**, Images show ectopic expression in a *mrc1a 125bp:GFP; kdrl:mCherry* double positive embryo that was injected with endothelial-specific *nrp1b:*ETV2, SOX7, Nr2f2. Red arrow points to GFP expression in an arterial EC. Dorsal aorta (DA); posterior cardinal vein (PCV). Scale bars represent 100 μm in **a**, **b** and **d**, and 250 μm in **c**.

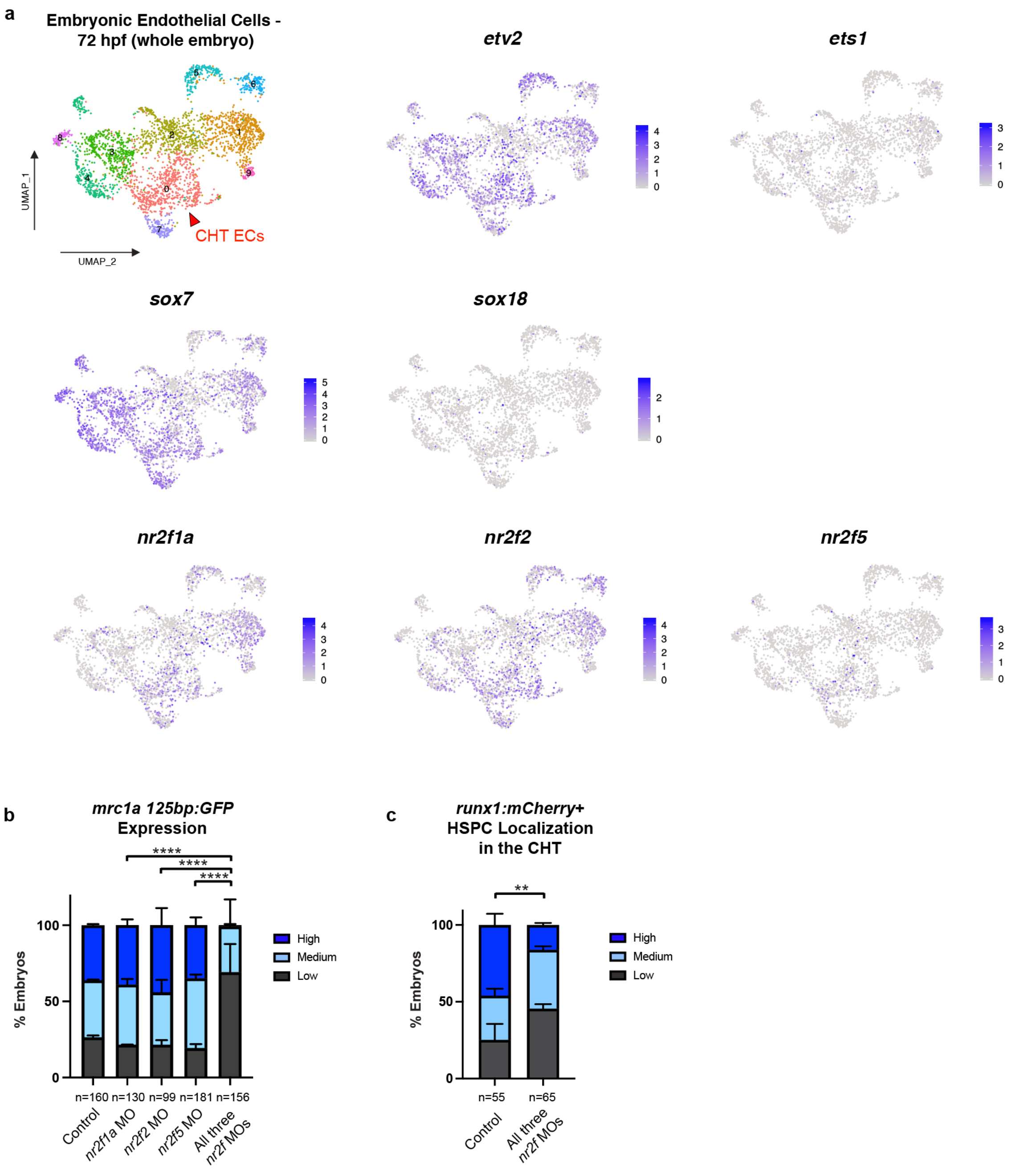


Extended Data Figure 9

**Extended Data Figure 9 | Redundancy in the transcription factor regulation of niche EC gene expression**. **a**, Uniform Manifold Approximation and Projection (UMAP) plots show cell clustering and gene expression from a single cell RNA-seq analysis of endothelial cells isolated from embryos at 72 hpf. Spectral scales report z-scores. **b**-**c,** Bar graphs report the quantification of *mrc1a 125bp:GFP* expression (**b**) and HSPC localization (**c**) in animals injected with the *nr2f1a, nr2f2, nr2f5* or control morpholinos. Chi squared test; $^{**}P<0.01$,$^{****}P<0.0001$.

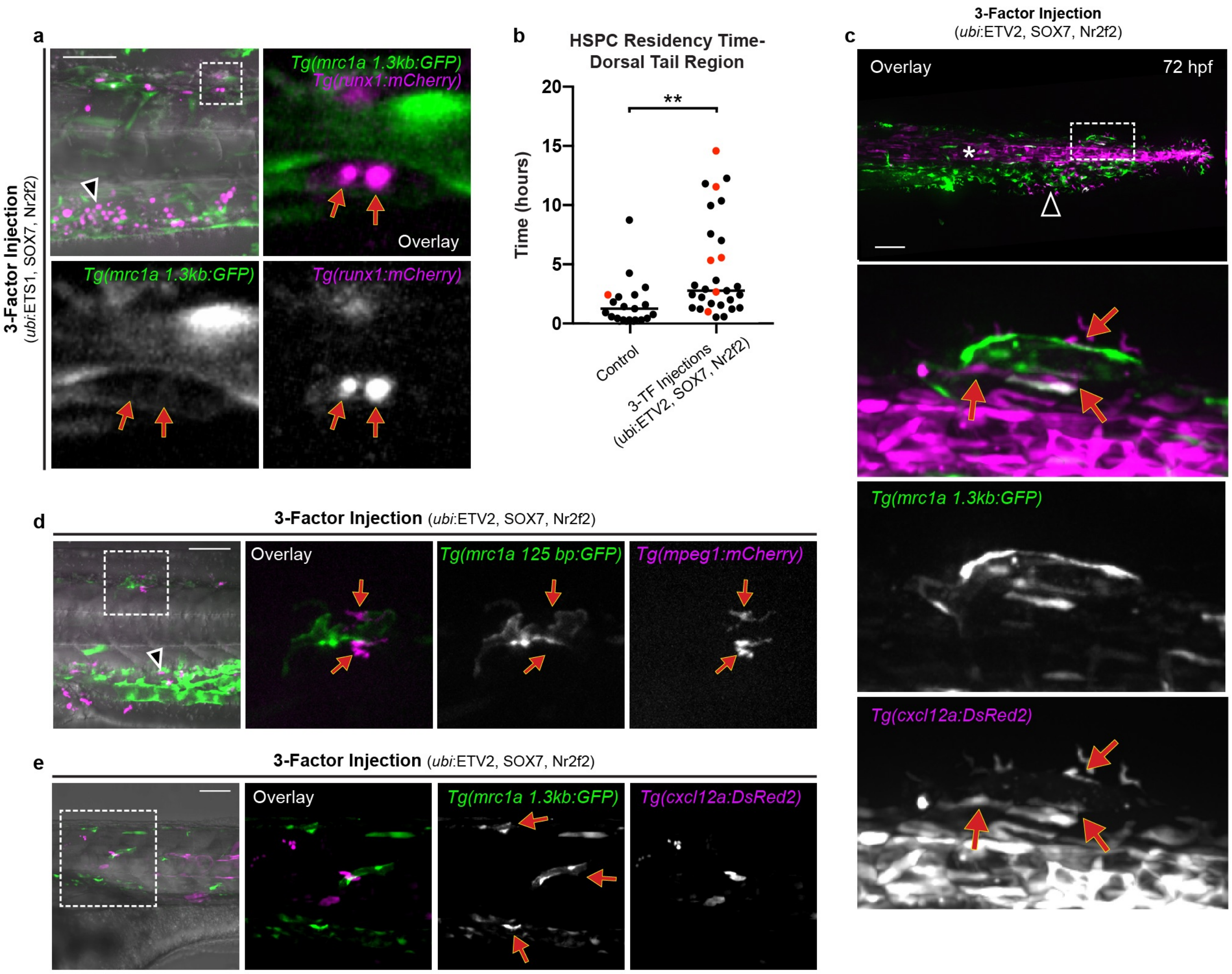


**Extended Data Figure 10**

**Extended Data Figure 10 | Niche endothelial gene expression recruits and supports HSPCs**. **a**, Image shows *runx1:mCherry*$^{+}$ HSPCs localized outside the CHT within a dorsal ectopic region of *mrc1a 1.3kb:GFP* expression in an embryo injected with a pool of *ubi:*ETS1, SOX7 and Nr2f2. Magnifications of boxed region are shown. Red arrows point to ectopic expression or localization while black arrowheads point to normal expression or localization in this and other panels in this figure. **b**, Graph reports lifetime measurements of HSPC residency outside of the CHT in control and 3-factor injected embryos. Red dots correspond to cells that divided. Mann-Whitney test; **$P<0.01$. **c**, ECs ectopically expressing *mrc1a 1.3kb:GFP* are associated with *cxcl12a:DsRed2*$^{+}$ stromal cells, similar to ECs in the CHT. Asterisk denotes notochord expression of *cxcl12a*:DsRed. Magnification of boxed region is shown. **d**, Images show ECs ectopically expressing *mrc1a 125bp:GFP* (boxed region) that are associated with *mpeg1:mCherry*$^{+}$ macrophages (red arrows), similar to ECs in the CHT (arrowhead). This data corresponds to Supplementary Video 7. **e**, Images show ECs (red arrows) in an anterior region of an embryo ectopically expressing *mrc1a 125bp:GFP* that are not associated with *cxcl12a:DsRed2*$^{+}$ stromal cells. Scale bars represent 100 μm.

**Supplementary Video 1 | Ectopic niche ECs arise independent of the endogenous CHT vasculature.** Video shows four different embryos at 22 hpf expressing *mrc1a 125bp:GFP* (top) or *kdrl:mCherry* (bottom) injected with control (left) or *ubi*:ETV2, SOX7, Nr2f2 plasmids (right). Red arrows point to ectopic vessels and black arrowheads point to the endogenous CHT niche. The duration of the time-lapse is 20 hours; time intervals are 45 minutes and video plays at 7 frames per second (fps).

**Supplementary Video 2 | Blood circulation through region of ectopic niche endothelial gene expression.** DIC time-lapse focusing up and down through a 72 hpf embryo shows healthy blood circulation through an ectopic patch of CHT-like vessels in the tail region of an embryo injected with *ubi:*ETV2, SOX7, Nr2f2 plasmids. The duration of the time-lapse is 40 seconds; time intervals are 150 milliseconds and video plays at 7 fps. The video corresponds to the embryo shown in Fig. 4c.

**Supplementary Video 3 | *mrc1a 125bp:GFP*$^+$ muscle cells change shape and integrate into the vasculature.** Video shows muscle-shaped cells ectopically expressing the *mrc1a 125bp:GFP* transgene (red arrows) that undergo morphological changes and integrate into the developing vasculature, in an embryo injected with *ubi:*ETV2, SOX7, Nr2f2 plasmids. The duration of the time-lapse is 20 hours; time intervals are 45 minutes and video plays at 7 fps.

**Supplementary Video 4 | *mrc1a 125bp:GFP*$^+$ muscle cells reorganize and change shape.** Video shows muscle-shaped cells within a somite that are ectopically expressing the *mrc1a 125bp:GFP* transgene (red arrow). In the video the cells undergo morphological changes that are uncharacteristic of normal muscle cells. The duration of the time-lapse is 12 hours; time intervals are 12 minutes and video plays at 7 fps. The video corresponds to the embryo shown in Extended Data Fig. 7d.

**Supplementary Video 5 | *mrc1a 125bp:GFP*$^+$ muscle cells become protrusive and migratory.** Video shows a muscle-shaped cell in the trunk of an embryo ectopically expressing the *mrc1a 125bp:GFP* transgene (red arrow). The animal is oriented with the posterior towards the bottom right. As the video plays, the cell develops protrusions and becomes migratory, processes that are uncharacteristic of normal muscle cells. One third of the way through the video a GFP+ cell dies or fragments into smaller pieces. The duration of the time-lapse is 6 hours; time intervals are 10 minutes and video plays at 7 fps.

**Supplementary Video 6 | Ectopic *mrc1a 125bp:GFP*$^+$ cells integrate into the vasculature.** Video shows an ectopic *mrc1a 1.3kb:GFP*$^+$ cell (green), in close proximity to a dorsal longitudinal anastomotic vessel (grayscale of *kdrl:mCherry* transgene is shown). As the video plays, the GFP$^+$ cell integrates and becomes part of the vasculature. The duration of the time-lapse is 20 hours; time intervals are 5 minutes and video plays at 7 fps.

**Supplementary Video 7 | Ectopic niche ECs are associated with macrophages.** Video shows ECs ectopically expressing the *mrc1a 125bp:GFP* transgene (green) that are associated with *mpeg1:mCherry*$^+$ macrophages (magenta; red arrow) in a 72 hpf embryo that had been injected with a pool of *ubi*:ETV2, SOX7 and Nr2f2 at the one cell stage. The duration of the time-lapse is 2 hours and 15 minutes; time intervals are 90 seconds and video plays at 7 fps. The video corresponds to the image shown in Extended Data Fig. 8j.

**Supplementary Video 8 | Initial recruitment of HSPC to region of ectopic niche endothelial expression.** Video shows a *runx1*$^+$ HSPC (magenta) initially lodging in a dorsal vessel that is ectopically expressing *mrc1a 1.3kb:GFP* (red arrow) in a 72 hpf embryo that had been injected with a pool of *ubi:*ETV2, SOX7 and Nr2f2 at the one cell-stage. Black arrowhead points to HSPC localization in the CHT. The duration of the time-lapse is 6.5 hours; time intervals are 2 minutes and the video plays at 10 fps. The video corresponds to data shown in Fig. 4d; images in Fig. 4d show still frame magnifications of the region in the video highlighted with the red arrow.

**Supplementary Video 9 | Proliferation of HSPCs and egress from ectopic region of niche endothelial gene expression.** Video shows *runx1*$^+$ HSPCs (magenta) localized to a vessel ectopically expressing *mrc1a 1.3kb:GFP* (red arrow) in a 72 hpf embryo that had been injected with a pool of *ubi*:ETV2, SOX7 and Nr2f2 at the one cell stage. HSPCs divide several times and migrate away into circulation. Black arrowhead points to HSPC localization in the CHT. The duration of the time-lapse is 2.6 hours; time intervals are 2 minutes and video plays at 10 fps.

**Supplementary Table 1 | CHT-enriched genes identified by tomo-seq**

| Gene | Full Gene Name | CHT Expression Confirmed by WISH |
|---|---|---|
| *abi3bp* | *ABI family, member 3 (NESH) binding protein* | Yes |
| *ACKR3* | *atypical chemokine receptor 3* | No* |
| *adam8a* | *ADAM metallopeptidase domain 8a* | Yes |
| *adra1d* | *adrenoceptor alpha 1D* | No* |
| *adrb3b* | *adrenoceptor beta 3b* | No* |
| *agrp* | *agouti related neuropeptide* | No* |
| *ANGPT4* | *angiopoietin 4* | No* |
| *ap1b1* | *adaptor-related protein complex 1, beta 1 subunit* | Yes |
| *aplnra* | *apelin receptor a* | Yes |
| *aqp7* | *aquaporin 7* | Yes |
| *atp1a1a.2* | *ATPase Na+/K+ transporting subunit alpha 1a, tandem duplicate 2* | n/a† |
| *atp1b1b* | *ATPase, Na+/K+ transporting, beta 1b* | n/a† |
| *ba1* | *ba1 globin* | n/a† |
| *blf* | *bloody fingers* | n/a† |
| *BX005069.4* | *leukocyte cell-derived chemotaxin-2-like* | n/a† |
| *BX323861.1* | *SLAM family member 9-like isoform X2* | n/a† |
| *C10H8orf4* | | n/a† |
| *ca15a* | *carbonic anhydrase XVa* | n/a† |
| *CABZ01049362.1* | *PREDICTED: GTPase IMAP family member 4-like [Danio rerio]. or 8-like* | n/a† |
| *CABZ01058863.1* | | n/a† |
| *CABZ01066695.1* | *PREDICTED: protein lyl-1-like isoform X1 [Danio rerio]./..lymphocytic leukemia protein* | n/a† |
| *ccdc88b* | *coiled-coil domain containing 88B* | n/a† |
| *ccr9a* | *chemokine (C-C motif) receptor 9a* | Yes |
| *CD209* | *CD209 molecule* | n/a† |
| *cd28* | *CD28 molecule* | Yes |
| *ceacam1* | *carcinoembryonic antigen-related cell adhesion molecule 1* | Yes |
| *ch25hl2* | *cholesterol 25-hydroxylase like 2* | n/a† |
| *cldn11a* | *claudin 11a* | Yes |
| *cldng* | *claudin g* | n/a† |
| *cltca* | *clathrin, heavy chain a (Hc)* | Yes |
| *cmklr1* | *chemokine-like receptor 1* | n/a† |
| *cndp2* | *carnosine dipeptidase 2* | Yes |
| *cnn1a* | *calponin 1, basic, smooth muscle, a* | Yes |
| *COL19A1* | *collagen, type XIX, alpha 1* | Yes |
| *col28a1* | *collagen, type 28, alpha 1* | n/a† |
| *coro1a* | *coronin, actin binding protein, 1A* | n/a† |
| *cpa5* | *carboxypeptidase A5* | n/a† |
| *CR381673.2* | *Natural killer cell receptor 2B4-like isorm x1/2 or SLAM family member7/9-like* | n/a† |
| *ctsh* | *cathepsin H* | Yes |
| *ctsla* | *cathepsin La* | Yes |
| *CU463790.1* | | Yes |
| *CU861664.1* | *PREDICTED: zinc finger protein 521-like [Danio rerio]* | n/a† |
| *CU915778.1* | *CU915778.1* | No* |
| *cyp24a1* | *cytochrome P450, family 24, subfamily A, polypeptide 1* | No* |
| *cysltr1* | *cysteinyl leukotriene receptor 1* | Yes |
| *dab2* | *Dab, mitogen-responsive phosphoprotein, homolog 2 (Drosophila)* | n/a† |
| *drl* | *draculin* | n/a† |
| *ela2* | *elastase 2* | Yes |
| *ENSDARG00000075833* | *lymphatic vessel endothelial hyaluronic receptor 1a/b* | Yes |
| *entpd2a.1* | *ectonucleoside triphosphate diphosphohydrolase 2a, tandem duplicate 1* | n/a† |
| *exoc3l2a* | *exocyst complex component 3-like 2a* | Yes |
| *f2r* | *coagulation factor II (thrombin) receptor* | Yes |
| *foxi3b* | *forkhead box I3b* | n/a† |
| *frem1b* | *Fras1 related extracellular matrix 1b* | Yes |
| *gcm2* | *glial cells missing homolog 2 (Drosophila)* | Yes |
| *glud1a* | *glutamate dehydrogenase 1a* | Yes |
| *glula* | *glutamate-ammonia ligase (glutamine synthase) a* | Yes |

| | | |
|---|---|---|
| *GMIP* | *GEM interacting protein* | n/a† |
| *gpr182* | *G protein-coupled receptor 182* | Yes |
| *grap2b* | *GRB2-related adaptor protein 2b* | n/a† |
| *gsto1* | *glutathione S-transferase omega 1* | Yes |
| *havcr1* | *hepatitis A virus cellular receptor 1* | n/a† |
| *hbaa1* | *hemoglobin, alpha adult 1* | n/a† |
| *hdr* | *hematopoietic death receptor* | n/a† |
| *hexb* | *hexosaminidase B (beta polypeptide)* | Yes |
| *hyal2a* | *hyaluronidase 2* | n/a† |
| *hyal2b* | *hyaluronidase 2* | Yes |
| *HYAL2* | *hyaluronidase 2* | Yes |
| *ifi30* | *interferon, gamma-inducible protein 30* | Yes |
| *il10ra* | *interleukin 10 receptor, alpha* | n/a† |
| *il13ra2* | *interleukin 13 receptor, alpha 2* | Yes |
| *il6r* | *interleukin 6 receptor* | Yes |
| *ITGAE* | *integrin, alpha E, tandem duplicate 1/2* | No* |
| *itgb2* | *integrin, beta 2* | Yes |
| *kcnj1a.3* | *potassium inwardly-rectifying channel, subfamily J, member 1a, tandem duplicate 3* | n/a† |
| *kcnj1a.5* | *potassium inwardly-rectifying channel, subfamily J, member 1a, tandem duplicate 5* | n/a† |
| *lamp2* | *lysosomal-associated membrane protein 2* | Yes |
| *lgals9l1* | *lectin, galactoside-binding, soluble, 9 (galectin 9)-like 1* | n/a† |
| *lgmn* | *legumain* | Yes |
| *lpar5a* | *lysophosphatidic acid receptor 5a* | n/a† |
| *mafbb* | *v-maf avian musculoaponeurotic fibrosarcoma oncogene homolog Bb* | n/a† |
| *man2b2* | *mannosidase, alpha, class 2B, member 2* | Yes |
| *marco* | *macrophage receptor with collagenous structure* | n/a† |
| *MCOLN2* | *mucolipin 2[WM2]* | n/a† |
| *mir142a* | *micro RNA 142a* | n/a† |
| *mmp13a* | *matrix metallopeptidase 13a* | n/a† |
| *MOV10L1* | *putative helicase Mov10l1 [Danio rerio]. 95% ident.* | n/a† |
| *mpx* | *myeloid-specific peroxidase* | n/a† |
| *mrc1a* | *mannose receptor, C type 1a* | Yes |
| *mrc1b* | *mannose receptor, C type 1b* | n/a† |
| *myh11a* | *myosin, heavy chain 11a, smooth muscle* | Yes |
| *myha* | *myosin, heavy chain a* | n/a† |
| *myo1f* | *myosin IF* | Yes |
| *ncf1* | *neutrophil cytosolic factor 1* | n/a† |
| *npl* | *N-acetylneuraminate pyruvate lyase (dihydrodipicolinate synthase)* | n/a† |
| *ostf1* | *osteoclast stimulating factor 1* | n/a† |
| *parvg* | *parvin, gamma* | Yes |
| *pdia2* | *protein disulfide isomerase family A, member 2* | n/a† |
| *PLCXD1* | *phosphatidylinositol-specific phospholipase C, X domain containing 1* | n/a† |
| *plek* | *pleckstrin* | n/a† |
| *polm* | *polymerase (DNA directed), mu* | n/a† |
| *prcp* | *prolylcarboxypeptidase (angiotensinase C)* | n/a† |
| *pxk* | *PX domain containing serine/threonine kinase* | No* |
| *rasal3* | *RAS protein activator like 3* | n/a† |
| *RNF223* | *ring finger protein 223* | n/a† |
| *s1pr4* | *sphingosine-1-phosphate receptor 4* | n/a† |
| *sele* | *selectin E* | Yes |
| *sepp1a* | *selenoprotein P* | Yes |
| *setx* | *senataxin* | Yes |
| *si:ch1073-429i10.1* | *si:ch1073-429i10.1* | Yes |
| *si:ch211-214p16.1* | *si:ch211-214p16.1* | No* |
| *si:ch211-214p16.2* | *si:ch211-214p16.2* | Yes |
| *si:ch211-250g4.3* | *PREDICTED: nesprin-1 isoform X4 [Danio rerio]* | n/a† |
| *si:ch211-284o19.8* | *si:ch211-284o19.8* | n/a† |
| *si:ch211-285f17.1* | *si:ch211-285f17.1* | n/a† |
| *si:ch211-67e16.2* | *cd28-like molecule* | n/a† |
| *si:ch73-248e21.7* | *si:ch73-248e21.7* | n/a† |
| *si:ch73-27e22.6* | *si:ch73-27e22.6* | n/a† |
| *si:dkey-102g19.3* | *si:dkey-102g19.3* | n/a† |

| | | |
|---|---|---|
| *si:dkey-188i13.7* | *interferon alpha inducible protein 46* | n/a† |
| *si:dkey-237j10.2* | *si:dkey-237j10.2* | n/a† |
| *si:dkey-33i11.4* | *si:dkey-33i11.4* | Yes |
| *si:dkey-69c1.1* | *si:dkey-69c1.1* | n/a† |
| *skap2* | *src kinase associated phosphoprotein 2* | n/a† |
| *sla1* | *src-like-adaptor 1* | n/a† |
| *slc16a9a* | *solute carrier family 16, member 9a* | No* |
| *slc4a11* | *solute carrier family 4, sodium borate transporter, member 11* | n/a† |
| *snx8a* | *sorting nexin 8a* | Yes |
| *srgn* | *serglycin* | Yes |
| *stab1* | *stabilin 1* | Yes |
| *stab2* | *stabilin 2* | Yes |
| *syk* | *spleen tyrosine kinase* | n/a† |
| *tagapb* | *T-cell activation RhoGTPase activating protein b* | n/a† |
| *tll1* | *tolloid-like 1* | Yes |
| *tmem106a* | *transmembrane protein 106a* | Yes |
| *tnfsf12* | *TNF superfamily member 12* | n/a† |
| *tnni1b* | *troponin I type 1b (skeletal, slow)* | Yes |
| *tubb1* | *tubulin, beta 1 class VI* | n/a† |
| *wasa* | *Wiskott-Aldrich syndrome (eczema-thrombocytopenia) a* | n/a† |
| *wasb* | *Wiskott-Aldrich syndrome (eczema-thrombocytopenia) b* | n/a† |
| *WIPF1* | *WAS/WASL interacting protein family, member 1a/b* | n/a† |
| *zgc:158446* | *complement factor b, like* | n/a† |
| *zgc:174945* | *zgc:174945* | n/a† |
| *zgc:198419* | *ferritin, heavy polypeptide-like 28* | Yes |

*No CHT expression was observed by WISH. †Did not attempt WISH.

**Supplementary Table 2 | CHT EC-enriched genes**

| Gene | Full Gene Name | CHT Expression Confirmed by WISH | Associated with CHT EC ATAC-seq Element* | Function |
|---|---|---|---|---|
| *adra1d* | *adrenoreceptor alpha 1D* | No† | Yes | G-protein coupled receptor; mitogenic response activation |
| *ap1b1* | *adaptor-related protein complex 1, beta 1 subunit* | Yes | Yes | Coated vesicle clathrin recruitment |
| *cldn11a* | *claudin 11a* | Yes | Yes | Tight junction strand component |
| *cltca* | *clathrin, heavy chain a (Hc)* | Yes | Yes | Major coated vesicle and coated pit component |
| *ctsh* | *cathepsin H* | Yes | No | Lysosomal cysteine proteinase |
| *ctsla* | *cathepsin La* | Yes | Yes | Lysosomal cysteine proteinase |
| *dab2* | *Dab, mitogen-responsive phosphoprotein, homolog 2 (Drosophila)* | n/a‡ | Yes | Mitogen-responsive phosphoprotein; clathrin-mediated endocytosis |
| *exoc3l2a* | *exocyst complex component 3-like 2a* | Yes | Yes | SNARE binding |
| *glula* | *glutamate-ammonia ligase (glutamine synthase) a* | Yes | Yes | Glutamine synthesis |
| *gpr182* | *G protein-coupled receptor 182* | Yes | Yes | G-protein coupled receptor; vasodilation |
| *hexb* | *hexosaminidase B (beta polypeptide)* | Yes | Yes | Degradation of N-acetyl hexosamine containing molecules |
| *hyal2a* | *hyaluronidase 2a* | n/a§ | Yes | Hyaluronan degradation |
| *hyal2b* | *hyaluronidase 2b* | Yes | Yes | Hyaluronan degradation |
| *ifi30* | *interferon, gamma-inducible protein 30* | Yes | Yes | Disulfide bond reduction, MHC class II-restricted antigen processing |
| *il13ra2* | *interleukin 13 receptor, alpha 2* | Yes | No | Interleukin 13 binding |
| *lgmn* | *legumain* | Yes | Yes | Hydrolysis of asparaginyl bonds |
| *lyve1b* | *lymphatic vessel endothelial hyaluronic receptor 1b* | Yes | Yes | Hyaluronan receptor |
| *man2b2* | *mannosidase, alpha, class 2B, member 2* | Yes | Yes | Mannose glycosylase |
| *mrc1a* | *mannose receptor, C type 1a* | Yes | Yes | Glycoprotein endocytosis |
| *npl* | *N-acetylneuraminate pyruvate lyase (dihydrodipicolinate synthase)* | n/a‡ | Yes | N-acetylneuraminic acid cleavage |
| *prcp* | *prolylcarboxypeptidase (angiotensinase C)* | n/a‡ | Yes | C-terminal proline linked amino acid cleavage |
| *pxk* | *PX domain containing serine/threonine kinase* | No | Yes | Synaptic transmission |
| *sele* | *selectin E* | Yes | Yes | Endothelial cell adhesion to blood leukocytes |
| *sepp1a* | *selenoprotein P* | Yes | No | Selenium binding |
| *slc16a9a* | *solute carrier family 16, member 9a* | No | Yes | Symporter activity |
| *snx8a* | *sorting nexin 8a* | Yes | Yes | Phosphatidylinositol binding |
| *stab1* | *stabilin 1* | Yes | Yes | Scavenger receptor activity |
| *stab2* | *stabilin 2* | Yes | Yes | Scavenger receptor activity, hyaluronan receptor |
| *tll1* | *tolloid-like 1* | Yes | Yes | Procollagen C-propeptide processing |

Table shows CHT EC genes identified by tomo-seq and tissue-specific RNA-seq. *Within 100 kb of TSS; some genes are associated with multiple elements. †No expression was observed by WISH. ‡Did not attempt WISH but CHT expression reported on zfin.org. §Did not attempt WISH.

**Supplementary Table 4 | *In vivo* screening of predicted enhancer elements**

| Type of Element | Gene Name | Genomic Coordinates of ATAC-seq Element[a] | Relative to TSS (kb) | Amplicon Size (bp) | Showed Predicted GFP Expression Pattern[b] | Element Contains Ets, SoxF and NHR Motifs |
|---|---|---|---|---|---|---|
| CHT EC Element | *ap1b1* | chr5:26463217-26463695 | 17 | 750 | Yes | Yes |
| | *cltca* | chr10:29,047,274-29,047,619 | 2.8 | 404 | Yes | Yes |
| | *dab2* | chr5:33,980,000-33,980,306 | -3.5 | 394 | Yes | Yes |
| | *exoc3l2a* | chr5:38359097-38359903 | 5.9 | 901 | Yes | Yes |
| | *glula* | chr2:19,458,704-19,459,047 | 4.8 | 446 | No | Yes |
| | *gpr182* | chr23:36701205-36701682 | -4.9 | 481 | Yes | Yes |
| | *gpr182* | chr23:36694073-36694476 | -2.8 | 398 | Yes | Yes |
| | *gpr182* | chr23:36696363-36696656 | 1.6 | 577 | Yes | Yes |
| | *lgmn* | chr13:36,448,465-36,448,818 | 2.9 | 414 | Yes | Yes |
| | *prcp* | chr15:10,400,588-10,400,868 | 23 | 334 | No | No[c] |
| | *sele* | chr20:34,010,027-34,010,326 | -9.7 | 398 | Yes | Yes |
| | *sele* | chr20:34,011,251-34,011,563 | -8.5 | 360 | Yes | Yes |
| | *snx8a* | chr3:42,090,805-42,091,062 | 5.5 | 395 | No | Yes |
| | *stab1* | chr22:10467346-10467937 | -2.8 | 874 | Yes | Yes |
| | *stab2* | chr4:9790795-9791116 | 4.3 | 422 | Yes | Yes |
| Pan-EC Element | *cdh5* | chr7:45457842-45458791 | 13 | 823 | Yes | Yes |
| | *clec14a* | chr17:10362325-10362844 | -3.1 | 455 | Yes | Yes |
| | *dll4* | chr20:28219013-28219619 | -55 | 452 | Yes | No[c] |
| | *fli1a* | chr18:47039842-47040466 | 47 | 800 | Yes | No[c] |
| | *lmo2* | chr18:36722030-36722527 | -3.6 | 367 | Yes | No[c] |
| | *nrp1b* | chr2:43535098-43535801 | -34 | 552 | Yes | Yes |

Table shows CHT EC-specific and pan-EC ATAC-seq elements that were fused to a minimal promoter and GFP and injected into one cell-stage zebrafish embryos. [a]Coordinates of MACS2 peak. [b]Expressed in CHT ECs for CHT EC elements and in vessels throughout the embryo for pan-EC elements. [c]Lacks NHR motif.

**Supplementary Table 5 | Genes highly enriched in CHT ECs versus head lymphatic ECs**

| | p_val | avg_logFC | pct.1 | pct.2 | p_val_adj |
|---|---|---|---|---|---|
| vcam1b | 2.18E-130 | 1.5050412 | 0.295 | 0.008 | 3.75E-126 |
| postnb | 4.06E-104 | 2.09928056 | 0.933 | 0.266 | 6.99E-100 |
| cfd | 3.82E-100 | 2.09891764 | 0.604 | 0.081 | 6.58E-96 |
| mrc1b | 7.80E-62 | 1.26023255 | 0.362 | 0.044 | 1.34E-57 |
| ifi30 | 2.99E-58 | 1.93882101 | 0.705 | 0.217 | 5.16E-54 |
| cyp1a | 1.09E-57 | 1.27880031 | 0.396 | 0.056 | 1.88E-53 |
| apoeb | 2.60E-56 | 1.13948574 | 0.329 | 0.039 | 4.48E-52 |
| slco2b1 | 2.31E-45 | 1.3134043 | 0.403 | 0.075 | 3.97E-41 |
| gpr182 | 4.29E-35 | 1.30290416 | 0.805 | 0.457 | 7.39E-31 |
| slit3 | 1.41E-32 | 0.99966049 | 0.383 | 0.09 | 2.42E-28 |
| snx8a | 2.27E-31 | 1.06913474 | 0.315 | 0.065 | 3.91E-27 |
| si.ch211.145 | 6.18E-30 | 1.24175318 | 0.725 | 0.378 | 1.06E-25 |
| slc3a2a | 1.46E-29 | 0.90195101 | 0.322 | 0.07 | 2.52E-25 |
| f13a1b | 3.93E-29 | 1.02693921 | 0.255 | 0.046 | 6.77E-25 |
| cplx2l | 4.20E-29 | 0.77751761 | 0.698 | 0.267 | 7.24E-25 |
| plvapb | 1.60E-25 | 0.73582562 | 0.919 | 0.609 | 2.75E-21 |
| mCherry | 1.67E-25 | 0.67608173 | 0.906 | 0.509 | 2.87E-21 |
| rgcc | 5.51E-25 | 0.91417728 | 0.523 | 0.188 | 9.49E-21 |
| slc43a1b | 2.13E-24 | 0.96912737 | 0.295 | 0.07 | 3.66E-20 |
| cebpb | 4.07E-24 | 1.13484095 | 0.409 | 0.132 | 7.00E-20 |
| aplnra | 1.04E-23 | 1.10743419 | 0.456 | 0.162 | 1.79E-19 |
| CR383676.1 | 4.67E-23 | 0.52085134 | 0.973 | 0.962 | 8.04E-19 |
| prcp | 5.87E-23 | 0.92014712 | 0.705 | 0.394 | 1.01E-18 |
| gstp1 | 2.55E-21 | 1.06867455 | 0.49 | 0.196 | 4.39E-17 |
| ctgfa | 3.31E-21 | 1.08499842 | 0.503 | 0.207 | 5.70E-17 |
| tmem176 | 5.02E-21 | 1.09849473 | 0.389 | 0.131 | 8.65E-17 |
| slc40a1 | 2.08E-19 | 1.04963187 | 0.396 | 0.145 | 3.58E-15 |
| sypl2a | 2.17E-19 | 0.85040214 | 0.765 | 0.512 | 3.73E-15 |
| cd81a | 3.74E-19 | 0.63350778 | 0.913 | 0.821 | 6.45E-15 |
| cox4i2 | 3.00E-18 | 0.94476235 | 0.302 | 0.09 | 5.17E-14 |
| arl4aa | 2.75E-17 | 0.94006877 | 0.436 | 0.186 | 4.74E-13 |
| selenbp1 | 4.14E-17 | 0.79509512 | 0.443 | 0.185 | 7.14E-13 |
| si.ch211.212 | 4.48E-17 | 0.96025024 | 0.544 | 0.263 | 7.71E-13 |
| ctsf | 1.64E-16 | 0.82323185 | 0.416 | 0.176 | 2.82E-12 |
| aplp2 | 1.69E-16 | 0.79851817 | 0.671 | 0.444 | 2.91E-12 |
| glud1a | 4.48E-16 | 0.91700097 | 0.537 | 0.28 | 7.71E-12 |
| creb3l3l | 4.57E-16 | 0.79298352 | 0.349 | 0.126 | 7.87E-12 |
| kirrel3l | 5.37E-16 | 0.79576482 | 0.463 | 0.202 | 9.24E-12 |
| frem1b | 5.45E-15 | 0.65276753 | 0.342 | 0.124 | 9.38E-11 |

| | | | | | |
|---|---|---|---|---|---|
| lmo2 | 1.14E-13 | 0.73839284 | 0.671 | 0.428 | 1.97E-09 |
| mafba | 3.01E-13 | 0.78147829 | 0.409 | 0.19 | 5.19E-09 |
| NC-002333.1 | 5.26E-13 | 0.61283393 | 0.832 | 0.697 | 9.06E-09 |
| pfkfb4b | 1.34E-12 | 0.65067785 | 0.302 | 0.116 | 2.31E-08 |
| sh3tc2 | 1.40E-12 | 0.7368562 | 0.383 | 0.174 | 2.40E-08 |
| sele | 2.83E-12 | 0.71507475 | 0.544 | 0.294 | 4.86E-08 |
| adcy7 | 3.56E-12 | 0.68474418 | 0.255 | 0.091 | 6.13E-08 |
| itm2bb | 5.62E-12 | 0.60693183 | 0.322 | 0.132 | 9.68E-08 |
| cxcl12a | 7.91E-12 | 1.02097412 | 0.517 | 0.287 | 1.36E-07 |
| nfe2l2a | 4.17E-11 | 0.64534152 | 0.376 | 0.175 | 7.18E-07 |
| NC-002333.4 | 5.25E-11 | 0.57667849 | 0.966 | 0.957 | 9.04E-07 |
| dap1b | 1.74E-10 | 0.58931482 | 0.523 | 0.314 | 3.00E-06 |
| sptlc2a | 1.46E-09 | 0.69935814 | 0.47 | 0.279 | 2.51E-05 |
| tspan4b | 4.70E-09 | 0.50277421 | 0.745 | 0.577 | 8.09E-05 |
| ctsk | 8.40E-09 | 0.59098275 | 0.282 | 0.127 | 0.00014472 |
| arhgdig | 1.79E-08 | 0.61012219 | 0.43 | 0.251 | 0.00030795 |
| fosab | 2.55E-08 | 0.49495873 | 0.846 | 0.8 | 0.00043855 |
| socs3a | 6.30E-08 | 0.56610115 | 0.517 | 0.32 | 0.00108405 |
| nr2f2 | 1.01E-07 | 0.54086981 | 0.483 | 0.302 | 0.00173123 |
| igf2b | 1.25E-07 | 0.29345929 | 0.43 | 0.236 | 0.0021557 |
| mdka | 2.02E-07 | 0.52272009 | 0.329 | 0.168 | 0.00348369 |
| fxyd6l | 2.50E-07 | 0.57025514 | 0.597 | 0.439 | 0.00430629 |
| arvcfb | 4.55E-07 | 0.6151207 | 0.262 | 0.127 | 0.00783973 |
| glrx | 1.05E-06 | 0.66550824 | 0.356 | 0.212 | 0.01812268 |
| eppk1 | 1.61E-06 | 0.41572371 | 0.302 | 0.157 | 0.02770418 |
| spns2 | 2.25E-06 | 0.49034806 | 0.409 | 0.255 | 0.03880785 |
| pdcd10a | 6.54E-06 | 0.63325722 | 0.289 | 0.165 | 0.11257852 |
| cotl1 | 8.94E-06 | 0.46044866 | 0.523 | 0.376 | 0.15390984 |
| foxo1a | 9.43E-06 | 0.46376427 | 0.302 | 0.169 | 0.1624564 |
| srgn | 9.86E-06 | 0.51260879 | 0.557 | 0.422 | 0.16972257 |
| atf3 | 1.32E-05 | 0.53387543 | 0.456 | 0.312 | 0.22644705 |
| atoh8 | 1.49E-05 | 0.43736127 | 0.255 | 0.134 | 0.25601762 |
| carhsp1 | 1.72E-05 | 0.44609223 | 0.389 | 0.256 | 0.29635763 |
| zgc.158343 | 1.83E-05 | 0.41378713 | 0.43 | 0.284 | 0.31510142 |
| tie1 | 3.41E-05 | 0.47469849 | 0.483 | 0.352 | 0.58796375 |
| CR318588.4 | 3.66E-05 | 0.30572062 | 0.866 | 0.764 | 0.63007722 |
| cav1 | 3.78E-05 | 0.25810291 | 0.584 | 0.402 | 0.65098699 |
| MDFIC | 5.29E-05 | 0.44498727 | 0.289 | 0.171 | 0.91030013 |
| gabarapb | 6.61E-05 | 0.38335703 | 0.584 | 0.461 | 1 |
| gadd45ba | 0.00010627 | 0.61613162 | 0.383 | 0.269 | 1 |
| klf11a | 0.00011116 | 0.47246032 | 0.309 | 0.19 | 1 |
| ednraa | 0.00012872 | 0.2821216 | 0.577 | 0.439 | 1 |

| | | | | | |
|---|---|---|---|---|---|
| cst3 | 0.00016627 | 0.40314002 | 0.362 | 0.248 | 1 |
| junbb | 0.00021583 | 0.37835247 | 0.597 | 0.5 | 1 |
| hif1al | 0.00022965 | 0.35097135 | 0.477 | 0.367 | 1 |
| ppp1r15a | 0.00025026 | 0.42112231 | 0.409 | 0.311 | 1 |
| fam129ba | 0.00037151 | 0.33286129 | 0.282 | 0.176 | 1 |
| antxr1c | 0.00043224 | 0.4234948 | 0.322 | 0.215 | 1 |
| wbp2nl | 0.00058622 | 0.52541004 | 0.255 | 0.162 | 1 |
| egr1 | 0.00082089 | 0.43273518 | 0.423 | 0.319 | 1 |
| afdna | 0.00095183 | 0.29296171 | 0.463 | 0.361 | 1 |
| fosl1a | 0.00115174 | 0.56840945 | 0.342 | 0.245 | 1 |
| cldn11a | 0.00122632 | 0.37114708 | 0.356 | 0.243 | 1 |
| AL929057.1 | 0.00135682 | 0.3080561 | 0.503 | 0.41 | 1 |
| txnipa | 0.00141179 | 0.39282414 | 0.409 | 0.312 | 1 |
| si.ch1073.29 | 0.00141489 | 0.42046256 | 0.255 | 0.168 | 1 |
| foxc1a | 0.0015422 | 0.41075596 | 0.423 | 0.314 | 1 |
| pmp22b | 0.00163391 | 0.30027848 | 0.55 | 0.453 | 1 |
| rab11a | 0.00177201 | 0.32073745 | 0.51 | 0.418 | 1 |
| arf2b | 0.00207173 | 0.32011369 | 0.477 | 0.377 | 1 |
| dspa | 0.0023146 | 0.28904343 | 0.329 | 0.234 | 1 |
| ier2b | 0.00249081 | 0.32304079 | 0.617 | 0.493 | 1 |
| ccdc85b | 0.00270545 | 0.40674321 | 0.362 | 0.275 | 1 |
| ptprb | 0.00271488 | 0.2642841 | 0.389 | 0.287 | 1 |
| fmnl3 | 0.00299415 | 0.31051483 | 0.477 | 0.392 | 1 |
| zgc.92066 | 0.00356946 | 0.41767579 | 0.638 | 0.599 | 1 |
| chmp2a | 0.00478999 | 0.39156411 | 0.362 | 0.269 | 1 |
| si.ch211.66e: | 0.00551295 | 0.26974094 | 0.309 | 0.224 | 1 |
| mbnl2 | 0.00606274 | 0.34652049 | 0.282 | 0.205 | 1 |
| tcima | 0.00707508 | 0.3469472 | 0.329 | 0.248 | 1 |
| fgd5a | 0.0073116 | 0.30294298 | 0.53 | 0.457 | 1 |
| mgst3b | 0.0075897 | 0.25460154 | 0.584 | 0.52 | 1 |
| fosb | 0.0078554 | 0.28043474 | 0.376 | 0.285 | 1 |
| capns1a | 0.01005763 | 0.27357761 | 0.282 | 0.206 | 1 |
| mcl1a | 0.01019623 | 0.34272388 | 0.463 | 0.392 | 1 |
| nccrp1 | 0.01030391 | 0.33916694 | 0.275 | 0.205 | 1 |
| chmp1b | 0.01093426 | 0.33988326 | 0.262 | 0.188 | 1 |
| midn | 0.0213744 | 0.38857182 | 0.409 | 0.355 | 1 |
| jam3b | 0.02162813 | 0.45849993 | 0.362 | 0.292 | 1 |
| lima1a | 0.02430426 | 0.28283817 | 0.342 | 0.268 | 1 |
| rps27l | 0.02455113 | 0.35521679 | 0.262 | 0.196 | 1 |
| cmip | 0.02470793 | 0.36383225 | 0.282 | 0.222 | 1 |
| klf2a | 0.03206854 | 0.26622252 | 0.389 | 0.327 | 1 |
| ak1 | 0.03306259 | 0.43558065 | 0.255 | 0.189 | 1 |

| | | | | | |
|---|---|---|---|---|---|
| rabgap1l | 0.03612582 | 0.4855517 | 0.268 | 0.207 | 1 |
| ier2a | 0.03634074 | 0.29031207 | 0.544 | 0.482 | 1 |
| cast | 0.0387489 | 0.26699972 | 0.302 | 0.239 | 1 |
| id3 | 0.03899849 | 0.29128838 | 0.403 | 0.35 | 1 |
| gabarapa | 0.04145586 | 0.34150085 | 0.268 | 0.212 | 1 |
| vps28 | 0.0443803 | 0.25469966 | 0.255 | 0.193 | 1 |
| si.dkey.42i9.[illegible] | 0.05375849 | 0.29720031 | 0.322 | 0.274 | 1 |
| ginm1 | 0.05543283 | 0.30813697 | 0.295 | 0.241 | 1 |
| tm9sf2 | 0.07300028 | 0.28485757 | 0.282 | 0.236 | 1 |
| pim1 | 0.07487454 | 0.2763142 | 0.483 | 0.456 | 1 |
| rps27.2 | 0.08333904 | 0.30355184 | 0.611 | 0.6 | 1 |
| fam43a | 0.08821304 | 0.30464162 | 0.356 | 0.317 | 1 |
| rab2a | 0.09741086 | 0.29199533 | 0.416 | 0.383 | 1 |
| romo1 | 0.12274659 | 0.31570211 | 0.396 | 0.376 | 1 |
| btg2 | 0.14381666 | 0.29059396 | 0.51 | 0.493 | 1 |
| caprin1b | 0.14461019 | 0.26589169 | 0.329 | 0.292 | 1 |
| lfng | 0.1466735 | 0.25032028 | 0.356 | 0.316 | 1 |
| sri | 0.15271421 | 0.30293128 | 0.416 | 0.392 | 1 |
| sdc4 | 0.18137997 | 0.2545621 | 0.362 | 0.335 | 1 |
| iqsec1b | 0.21625231 | 0.27442777 | 0.262 | 0.23 | 1 |
| vta1 | 0.28421405 | 0.31193127 | 0.268 | 0.243 | 1 |
| rsrp1 | 0.38481374 | 0.26735815 | 0.477 | 0.497 | 1 |
| si.dkey.177p[illegible] | 0.40525573 | 0.27348436 | 0.322 | 0.309 | 1 |
| dusp1 | 0.55715268 | 0.34338202 | 0.309 | 0.308 | 1 |

**Supplementary Table 6 | Transcription factor expression in CHT ECs**

| Transcription Factor | Family | FPKM | Associated with CHT EC ATAC-seq Element Containing Ets, Sox and NHR Sites* | Genomic Coordinates of Representative Element |
|---|---|---|---|---|
| *fli1a* | Ets | 480.4 | Yes | chr18:46966409-46966698 |
| *etv2* | Ets | 192.3 | Yes | chr16:44782409-44782895 |
| *ets1* | Ets | 183 | Yes | chr18:46883643-46884100 |
| *sox18* | SoxF | 206.4 | Yes | chr23:8886011-8886744 |
| *sox7* | SoxF | 125.1 | Yes | chr20:19158376-19158663 |
| *nr2f2* | NHR | 84.6 | Yes | chr18:23728906-23729747 |
| *rxraa* | NHR | 45.9 | Yes | chr21:16411020-16411531 |

Table shows FPKM expression values in CHT ECs for highly expressed members of the Ets, Sox and NHR transcription factor families.
*Within 100 kb of TSS; some genes are associated with multiple elements.

**Supplementary Table 7 | Transcription factor expression in mouse hematopoietic niche**

| Transcription Factor | Family | Mouse E14-E15 Liver EC FPKM | Mouse E16-E17 Liver EC FPKM | Mouse Adult Bone Marrow EC FPKM |
|---|---|---|---|---|
| *Ets1* | Ets | 218.4666 | 251.9493 | 153.2657 |
| *Erg* | Ets | 46.64156 | 78.53131 | 45.14673 |
| *Elk4* | Ets | 9.369453 | 11.4226 | 22.83457 |
| *Elk1* | Ets | 7.003965 | 9.08418 | 6.8779 |
| *Etv1* | Ets | 2.203135 | 3.10327 | 1.488542 |
| *Etv2* | Ets | 0.235977 | 0 | 0 |
| *Sox18* | SoxF | 127.1509 | 262.44 | 130.1783 |
| *Sox7* | SoxF | 49.94503 | 46.9365 | 19.80563 |
| *Sox17* | SoxF | 33.01219 | 68.37438 | 90.24645 |
| *Sox11* | SoxF | 12.01665 | 11.1584 | 0.67509 |
| *Sox12* | SoxF | 11.81507 | 21.5267 | 0.556478 |
| *Sox6* | SoxF | 1.741399 | 1.158524 | 0.51182 |
| *Sox5* | SoxF | 0.193041 | 0.289841 | 0.437005 |
| *Sox9* | SoxF | 0.119527 | 0.072563 | 0 |
| *Nr2f2* | NHR | 58.97832 | 103.5558 | 63.17458 |
| *Rxra* | NHR | 23.98264 | 33.0942 | 22.08392 |
| *Rara* | NHR | 19.29841 | 27.37294 | 13.93433 |
| *Nr4a2* | NHR | 10.13413 | 3.130986 | 30.30394 |
| *Esrrb* | NHR | 6.219864 | 7.884516 | 0.586381 |
| *Rora* | NHR | 1.219604 | 1.086872 | 5.922048 |

**Supplementary Table 8 | Primers used for WISH probe synthesis**

| Category | Gene | Forward* | Reverse† |
|---|---|---|---|
| CHT EC enriched | *adra1d* | GCTCCATAGTATCGTCTGAACC | AAACCATTGCCATTTTGCCA |
| CHT EC enriched | *ap1b1* | GGGAGTTCTTCGGGTGACTG | GCTTGCAACAAAAAGCGCAG |
| CHT EC enriched | *cldn11a* | TGTGTGATCTCAACTGCGCT | GGTGCAATCTAGTCTGATCGGT |
| CHT EC enriched | *cltca* | CCAGCAAACCCCATGGATCT | AACCGAGTACAGGACACACG |
| CHT EC enriched | *ctsh* | CGACTGGAGAACCAAGGGAC | TGGAGGCTAATCGAGTGTGC |
| CHT EC enriched | *ctsla* | CCATGCAACAGAGGAAGGGT | TACTGGGCGGGTCTCCTTTA |
| CHT EC enriched | *exoc3l2a* | AAGTTCCGCAGGATGGACTG | TCGCTTGTGTGATCAAGTATGAC |
| CHT EC enriched | *glula* | AGTTATGCCAGCTCAGTGGG | GGCCTCCCCAAGAAACCATT |
| CHT EC enriched | *gpr182* | CTTCCCACAGCAGCACAAAC | GAAAGTTGTTGTTGAAGTGAACG |
| CHT EC enriched | *hexb* | GAATTTGCTCGCATGAGGGG | CGGCAGTGGCCAACAAATAG |
| CHT EC enriched | *hyal2b* | ATGGAGGTCTACCACAGGCT | AGTGCAGGTATGTGTCCGTG |
| CHT EC enriched | *ifi30* | TTCGGCTTTAACCTGTGCGT | CCTGACGCGAGTAGTGTTGT |
| CHT EC enriched | *il13ra2* | AGTTAGAATGGGCGCCACC | GGCAAGACCACTGGCATTTG |
| CHT EC enriched | *lgmn* | AACTTGAGCCACCGAGGATTT | CCCTAACTCCAGCACACACT |
| CHT EC enriched | *lyve1b* | GCTACAGTCTGCGTAGCAT | TGGAAGCAGCTCTAAGTGACAG |
| CHT EC enriched | *man2b2* | TACCCAATGGTTCGAGTGGC | GCTTAGGTGATCAATTTTGGGACA |
| CHT EC enriched | *mrc1a* | GTGTCCCCTCATCAATGCCA | ACGGCATTCCACAAACCAGA |
| CHT EC enriched | *sele* | TGCCCAGCCCTTGATAATCT | ACCCAAACTGACTTTATATGTGC |
| CHT EC enriched | *sepp1a* | AGGCAGCACTGGACTTTAGC | AGGTACAAATGCAAGTACAACACTG |
| CHT EC enriched | *snx8a* | ACAAAGAGATCTGCATTCCAAGC | AGCCTGTCAGCTCACTTTATT |
| CHT EC enriched | *stab1* | AAGGCGTACTATGTCCTCAGGC | CGCCGTTCTATAATGCACCG |
| CHT EC enriched | *stab2* | TTGTGGATTACGGGGTTCGG | AAAGAGAGCTGCACCGACT |
| CHT EC enriched | *tll1* | GAGCTTTACTCTGCTGGCGA | ACAAATGATGTCTGTCTCCGCT |
| CHT enriched | *abi3bp* | CTGTTTTCCCCCACCAGTGA | CAAAGGATTGGCAGGGACCA |
| CHT enriched | *ackr3* | TGGGATTTATTTGTAACTACACGGA | TTTTAAGCACATTTCTGAAGCACA |
| CHT enriched | *adam8a* | CCAGGAAGCGCAAAGAACAG | ACATTAGGCGGGCAAAACAAA |
| CHT enriched | *adrb3b* | GCAGCAAACGACTGCTACAA | CCCACTTCGCTGCTCTTTAC |
| CHT enriched | *agrp* | TCATCCACACCTGAGACGCA | ACACCTTAAAACCGCAGCC |
| CHT enriched | *angpt4* | ATCCGACTGCTGGAATGGAC | GCTTTGAGGAGCTTAAGAGGC |
| CHT enriched | *aplnra* | GTGCTGGTCAACATGTACGC | CGTCACTTTTCACCCCCAGA |
| CHT enriched | *aqp7* | TCCACTGGGAAAAGCTGGAAT | TTTCAGATGCAGCACAGGCA |
| CHT enriched | *bnip3lb* | ATGGGGCTGACGGATACC | GCACAGGAAACGCACATGAT |
| CHT enriched | *ccr9a* | TTGTCCAGACTACCAAGGCG | TTACTTCACTGCCAGTCGGC |
| CHT enriched | *cd28* | ATCCAAACTGAGGCCGGAAG | AGAAAATACAGTGCATACATGTCAA |
| CHT enriched | *ceacam1* | GGCCCAAGCATGGCAGAAAC | CCTACAAGCCTCATTCAGACAGT |
| CHT enriched | *cndp2* | ACATGGGACATGGAGCGAAG | ACACTAGAAAACCGATCGTGTCA |
| CHT enriched | *cnn1a* | GACTCTCTGCGGATGTCAGG | GGTCATGCCCTTTTGGCTTG |
| CHT enriched | *col19a1* | CATGTCCACCCCTGAAGCTG | GGGTTCTGTTGTGGAGTGCT |
| CHT enriched | *cu463790.1* | GGCGTCTCTTTTTCTGCTGC | TGACGCTTAAACAGAGCGGT |
| CHT enriched | *cu915778.1* | CCCTAGTGTCCGAGGTCTCA | TTTCCCCTGTGTGGATGAGC |
| CHT enriched | *cyp24a1* | GATACCGTGCTGGGCGATTA | CCACCACTCACTCATTCAGACA |

| | | | |
|---|---|---|---|
| CHT enriched | *cysltr1* | TCCCGGTGCAAAATCTGAGG | AGTCATGCACAAAATCTGCGG |
| CHT enriched | *ela2* | GTTTATTGCTGGCGCCTACG | TTCTTGGGGTAGTTGCAGCC |
| CHT enriched | *f2r* | GCTGCCGAACAACGAAACAT | TAGGACGCGTCATTGTGCTT |
| CHT enriched | *frem1b* | AGTACACTCCGGACCCAAGA | CACCAGAAAGAATGTCACCGT |
| CHT enriched | *gcm2* | TCCAGAGCGATTCAGCATCA | CAGTCCCTCAGTATTCCCCG |
| CHT enriched | *glud1a* | AGTCTCCTACTTCGAGTGGCT | ACGCCTGAGATTCATCCTGC |
| CHT enriched | *il6r* | AACTGTTCTTTCTCCCGGTCCC | CCTCTGGCTGAACAGGAAGG |
| CHT enriched | *itgae* | ACTGGTCAACCACCTCCTCT | ACACAATCAGGCAAGGTCTC |
| CHT enriched | *itgb2* | TGCCTTTCAAAGTGGACCGT | ACCAGTCACACCAGCCATTC |
| CHT enriched | *lamp2* | AGCCTGTTCCTGGACCATTG | AGCTACAACCATTGAGGGCT |
| CHT enriched | *myh11a* | GGTTCGCCAGAAGGACAAGA | AGCATCCAAAAGTACTCGGTGT |
| CHT enriched | *myo1f* | AAGCTGTCATCAAAGCCGGA | TTCTCGACCTGTCAGCTGTT |
| CHT enriched | *parvg* | TGAAAGCCCTGAACGAGACC | CGTCAGCATCCAAACGCAAT |
| CHT enriched | *setx* | AGGAGTTTGGCTTCGACCAG | GTGACGCTGGAATATCCCGT |
| CHT enriched | *si:ch1073-429i10.1* | TCGCTCTGATGCTCAGCTTG | CACTCGGCGACAGTATTCCC |
| CHT enriched | *si:ch211-214p16.1* | TACACATTTTCTGCCCCACTGA | AATGGGGCAAGAGTCCATCT |
| CHT enriched | *si:ch211-214p16.2* | CTCACCCTCGGTCCAGAACT | ACAGACACACTTGCCAGTCA |
| CHT enriched | *si:dkey-33i11.4* | ACAGCCATCAGTTCCTCTGC | AGCTTTGCATCCCCATCACT |
| CHT enriched | *srgn* | GGAAGCCACTCCTGATACGG | GTACAACATTTACTTGCTGTCCA |
| CHT enriched | *tmem106a* | GGTCACGCACCAAATGAACC | AACAGTTCTGATTGGATTTTGCTCA |
| CHT enriched | *tnni1b* | TCTGCATCTCGCAAGCTGAT | CATGTGTAGTGCAGACAGAACA |
| CHT enriched | *zgc:198419* | AGAACTACGACAGCGACTGC | GGTTTTGGATAAGAGCTGTGTCA |
| Transcription Factor | *ets1* | ACAGACTCTGTACGTTTGAATGCGT | GTCCAGACTTTACTCGTCCGTGTC |
| Transcription Factor | *etv2* | TATGACTGCAGTGGTGAAGACC | CTTTCCCGCCGTTTTGTGAA |
| Transcription Factor | *fli1a* | CAGACCCGTCTCTGTGGTC | CCAGTATGGGGTTGTGGGAC |
| Transcription Factor | *nr2f2* | ACCCCCGAACAACAATAACA | AGAGGGCAAGCGCAGTAATA |
| Transcription Factor | *sox7* | TATAGCCCTTCGTTCCCCCA | ACCGAAACCGGCTAAACTGA |
| Transcription Factor | *sox18* | TCCTTGGACGCTGTGGACCAAC | TCAAAGCGCTGCTTTCCTCGC |

*The T3 sequence CATTAACCCTCATAAAGGGAA was added to the 5' end of each forward primer.

†The T7 sequence TAATACGACTCACTATAGGG was added to the 5' end of each reverse primer.

**Supplementary Table 9 | Primers used to clone promoter and enhancer elements**

| Type of Element | Gene | Genomic Coordinates of ATAC-seq Element | Amplicon Size (bp) | Forward | Reverse |
|---|---|---|---|---|---|
| 5' upstream of TSS | *mrc1a* (1.3 kb) | chr7:65,468,213-65,469,565 | 1353 | CTTTTGGCCATTACTGCCG | TTCTGTCTTTTAATCAGCAATCC |
| CHT EC element | *mrc1a* (125 bp) | chr7:65469086-65469210 | 125 | GCTCTCAGTTCCTGGTATTTTTCT | TGAAGCTTGTACCTTTCATTTCC |
| 5' upstream of TSS | *sele* (5.3 kb) | chr20:34,001,481-34,006,781 | 5301 | TCGTTACTGCACTTGAAAGCGT | TATCAGTGATGTTCTGCAGTGGTC |
| CHT EC element | *sele* (158 bp) | chr20:34004805-34004962 | 158 | CCATGAAACTGGGAAGATGAA | CAGGAAGAAATAATGGCAAAAA |
| CHT EC element | *ap1b1* | chr5:26463217-26463695 | 750 | GAAGCTCTCCAGCAGCTCA | CATTTCCACCAGCTGTCTGAT |
| CHT EC element | *cltca* | chr10:29,047,274-29,047,619 | 404 | GCTGTCAGCACATTCTTTTCC | CCCTGCTGATCACACATGAC |
| CHT EC element | *dab2* | chr5:33,980,000-33,980,306 | 394 | ACTGCTCCTCACCAATCGTC | TGCACTAAATCTGTGCCAAGTC |
| CHT EC element | *exoc3l2a* | chr5:38359097-38359903 | 901 | TTTATATAATCGGAAGGAACCTTTTT | TCCTGTCAGCTGTTTTCATCC |
| CHT EC element | *glula* | chr2:19,458,704-19,459,047 | 446 | GGCAAAATGCTTAGATGCAGA | TGCGAGGAGGACATAAACAA |
| CHT EC element | *gpr182* | chr23:36701205-36701682 | 481 | TAGCCTTGTGCAATGCTTGT | TGCTGAATTCAAAAGCCACTT |
| CHT EC element | *gpr182* | chr23:36694073-36694476 | 398 | CACTTCTGGTACCAAATGATCAAC | GAGGGTTAAACGTGGCCTTA |
| CHT EC element | *gpr182* | chr23:36696363-36696656 | 577 | GCGGCAAACTTTTTGAGTGT | GCCAGCCTCAAAGTTTGTTCT |
| CHT EC element | *lgmn* | chr13:36,448,465-36,448,818 | 414 | CGCGTGATGAGGATCTGATT | GGTGTTGAAAGGTGATGCTG |
| CHT EC element | *prcp* | chr15:10,400,588-10,400,868 | 334 | AAAATTAAGAGCGGGCAGACT | TGGAAACAACAACAGCCTGA |
| CHT EC element | *sele* | chr20:34,010,027-34,010,326 | 398 | AAAGCACTTGATTGAGAATTGC | TGTTTGGTTCAGTTACACGTTTT |
| CHT EC element | *sele* | chr20:34,011,251-34,011,563 | 360 | CAGTTTCCCAAGCTTCAAGG | TGTGATTACACATTCCCACACAT |
| CHT EC element | *snx8a* | chr3:42,090,805-42,091,062 | 395 | AATGGTTGCAGCATTGTGTT | GCTTTTGTTTGGTGATGTGC |
| CHT EC element | *stab1* | chr22:10467346-10467937 | 874 | GTTACCTGGCAACCACCAAC | TGGTCAGAATAAGCACGTTTCA |
| CHT EC element | *stab2* | chr4:9790795-9791116 | 422 | ACGTTAACAAAGGCGATGTTTT | TCTAAACAATTTTTAAGGTAAACCAAA |
| Pan-EC element | *cdh5* | chr7:45457842-45458791 | 823 | TGACAGGACTCATCAGCACG | AATAGTCTCTGGTCTGCTGTTAAA |
| Pan-EC element | *clec14a* | chr17:10362325-10362844 | 455 | TGGGAAAAATACCAGGAAGCGT | AAGCAGCGAGCTCTCATAATAAA |
| Pan-EC element | *dll4* | chr20:28219013-28219619 | 452 | AGATCAATGAGAGCGAGGCG | GGAGCAGATGAGGTTAAGTCCT |
| Pan-EC element | *fli1a* | chr18:47039842-47040466 | 800 | CGGACAGTAATGTCTGGATGG | CCACAACTCCATACTGGGAAA |
| Pan-EC element | *lmo2* | chr18:36722030-36722527 | 367 | TCATCATGGCCAACAGAATG | GTGCAGGAAATGAGCACAGA |
| Pan-EC element | *nrp1b* | chr2:43535098-43535801 | 552 | TGACTCAACCAATCAATCAGCCT | TAGCAAAGCTCTCAGGCCC |

**Supplementary Table 10 | Sequences and primers for mutational variants of the 125 bp *mrc1a* and 158 bp *sele* enhancer elements**

| Gene | Fragment Name | Total Fragment Sequence | Forward Primer | Reverse Primer |
|---|---|---|---|---|
| *mrc1a* | Wild-type | CCATGAAACTGGGAAGATGAAAGCATTAGTTGAATTGTTACTGGCAACATCTTCTCTGTAATGCCCCCTGTGACCCATATTGTCTCGCTCTTTCCTTTATAAACAGAGCTGTAGATATCCACAGGAAATGGGGGTGTTTTTGCCATTATTTCTTCCTG | TGAAGCTTGTACCTTTCATTTCCTTTTTGCTGAGCTTTATTTTCTCTAGAATTGCCATTGTGTTTCCATTCTAG | GCTCTCAGTTCCTGGTATTTTTCTTTCAGCTGAAAAAAAAATGCTGATTTGCTAGAATGGAAACACAATGGCAAT |
| *mrc1a* | Ets mutant | TGAAGCTTGTACCTTTCATTTaaaTTTTGCTGAGCTTTATTTTCTCTAGAATTGCCATTGTGTTTCCATTCTAGCAAATCAGCATTTTTTTTTCAGCTGAAAGAAAAATACCAtttACTGAGAGC | TGAAGCTTGTACCTTTCATTTaaaTTTTGCTGAGCTTTATTTTCTCTAGAATTGCCATTGTGTTTCCATTCTAG | GCTCTCAGTaaaTGGTATTTTTCTTTCAGCTGAAAAAAAAATGCTGATTTGCTAGAATGGAAACACAATGGCAAT |
| *mrc1a* | Sox mutant | TGAAGCTTGTACCTTTCATTTCCTTTTTGCTGAGCgggcggggaTCTAGAATTGCacggtgGTTTCCATTCTAGCAAATCAGCcggggggTTTCAGCTGAAAGAAAAATACCAGGAACTGAGAGC | TGAAGCTTGTACCTTTCATTTCCTTTTTGCTGAGCgggcggggaTCTAGAATTGCacggtgGTTTCCATTCTAG | GCTCTCAGTTCCTGGTATTTTTCTTTCAGCTGAAAccccccgGCTGATTTGCTAGAATGGAAACcaccgtGCAAT |
| *mrc1a* | NHR mutant | attAGCagatTtaaTTTCATTTCCTTTTTGCattaaTTTATTTTCTCTAGAATTGCCATTGTGTTTCCATTCTAGCAAATCAGCATTTTTTTTTCAGCTGAAAGAAAAATACCAGGAACTGAGAGC | attAGCagatTtaaTTTCATTTCCTTTTTGCattaaTTTATTTTCTCTAGAATTGCCATTGTGTTTCCATTCTA | GCTCTCAGTTCCTGGTATTTTTCTTTCAGCTGAAAAAAAAATGCTGATTTGCTAGAATGGAAACACAATGGCAAT |
| *mrc1a* | Control mutant | TGAAGCTTGTACCTTTCATTTCCTTTTTGCTGAGCTTTATTTTCTCTAGAATTGCCATTGTGTTTCCATTCcgGCAAATCAGCATTTTTTTTTCAGCTGAccGAAAAATACCAGGAACTGAGAGC | TGAAGCTTGTACCTTTCATTTCCTTTTTGCTGAGCTTTATTTTCTCTAGAATTGCCATTGTGTTTCCATTCTAG | GCTCTCAGTTCCTGGTATTTTTCggTCAGCTGAAAAAAAAATGCTGATTTGCcgGAATGGAAACACAATGGCAAT |
| *sele* | Wild-type | CCATGAAACTGGGAAGATGAAAGCATTAGTTGAATTGTTACTGGCAACATCTTCTCTGTAATGCCCCCTGTGACCCATATTGTCTCGCTCTTTCCTTTATAAACAGAGCTGTAGATATCCACAGGAAATGGGGGTGTTTTTGCCATTATTTCTTCCTG | CCATGAAACTGGGAAGATGAAAGCATTAGTTGAATTGTTACTGGCAACATCTTCTCTGTAATGCCCCCTGTGACCCATATTGTCTCGCTCT | CAGGAAGAAATAATGGCAAAAACACCCCCATTTCCTGTGGATATCTACAGCTCTGTTTATAAAGGAAAGAGCGAGACAATATGGGTCACAG |
| *sele* | Ets mutant | CCATGAAACTGGGAAGATGAAAGCATTAGTTGAATTGTTACTGGCAACATCTTCTCTGTAATGCCCCCTGTGACCCATATTGTCTCGCTCTTTaaaTTATAAACAGAGCTGTAGATATCCACAtttAATGGGGGTGTTTTTGCCATTATTTCTaaaTG | CCATGAAACTGGGAAGATGAAAGCATTAGTTGAATTGTTACTGGCAACATCTTCTCTGTAATGCCCCCTGTGACCCATATTGTCTCGCTCT | CAtttAGAAATAATGGCAAAAACACCCCCATTaaaTGTGGATATCTACAGCTCTGTTTATAAtttAAAGAGCGAGACAATATGGGTCACAG |
| *sele* | Sox mutant | CCATGAAACTGGGAAGATGAAAGCATTAGTTGAAggtggACTGGCAACATCTTCTCTGTAATGCCCCCTGTGACCCATAggtgaTCGCTCTTTCCTTTATAAACAGAGCTGTAGATATCCACAGGAAATGGGGGTGTTTTTGCCATTATTTCTTCCTG | CCATGAAACTGGGAAGATGAAAGCATTAGTTGAAggtggACTGGCAACATCTTCTCTGTAATGCCCCCTGTGACCCATAggtgaTCGCTCT | CAGGAAGtttattaGGCAAAAACACCCCCATTTCCTGTGGATATCTACAGCTCTGTTTATAAAGGAAAGAGCGAtcaccTATGGGTCACAG |
| *sele* | NHR mutant | CCATGAAACTGGGAAGATGAAAGCATTAGTTGAATTGTTACTGGCAACATCTTCTCTGTAATGCCCCCTGattaaCATATTGTCTCGCTCTTTCCTTTATAAACAGAGCTGTAGATATCCACAGGAAATGGGGGTGTTTTTGCCATTATTTCTTCCTG | CCATGAAACTGGGAAGATGAAAGCATTAGTTGAATTGTTACTGGCAACATCTTCTCTGTAATGCCCCCTGattaaCATATTGTCTCGCTCT | CAGGAAGAAATAATGGCAAAAACACCCCCATTTCCTGTGGATATCTACAGCTCTGTTTATAAAGGAAAGAGCGAGACAATATGttaatCAG |
| *sele* | Control mutant | CCATGAAACTGGGAAtcTGAAAGCATTAGTTGAATTGTTACTGGCAACATCTTCTCTGTAATGCCCCCTGTGACCCATATTGTCTCGCTCTTTCCTTTATAAACAGAGagGTAGATATCCACAGGAAATGGGGGacTTTTTGCCATTATTTCTTCCTG | CCATGAAACTGGGAAtcTGAAAGCATTAGTTGAATTGTTACTGGCAACATCTTCTCTGTAATGCCCCCTGTGACCCATATTGTCTCGCTCT | CAGGAAGAAATAATGGCAAAAAgtCCCCCATTTCCTGTGGATATCTACctCTCTGTTTATAAAGGAAAGAGCGAGACAATATGGGTCACAG |

Lowercase letters indicate base pair changes used to disrupt transcription factor binding motifs.

**Supplementary Table 11 | Primers used for cloning and EMSA probe synthesis**

| Category | Primer Name | Forward | Reverse | Comment |
|---|---|---|---|---|
| Cloning | *Nr2f2* | CGGGATCCatggca atggtagtca gcacg | CCGGAATTCCGGttgaattgccatatatggc | |
| Probe synthesis | *mrc1a* site 1 wild-type | tttaTGAAGCTTGTACCTTTCATTTCCTTTTTG | CAAAAAGGAAATGAAAGGTACAAGCTTCAtaaa | |
| Probe synthesis | *mrc1a* site 1 mutation | TTTAattAGCagatTtaaTTTCATTTCCTTTTTG | CAAAAAGGAAATGAAAttaAatctGCTaatTAAA | 1st NHR site mutated like *in vivo* GFP reporter experiment |
| Probe synthesis | *mrc1a* site 2 wild-type | TTCATTTCCTTTTTGCTGAGCTTTATTTTC | GAAAATAAAGCTCAGCAAAAAGGAAATGAA | |
| Probe synthesis | *mrc1a* site 2 mutation | TTCATTTCCTTTTTGCattaaTTTATTTTC | GAAAATAAAttaatGCAAAAAGGAAATGAA | 2nd NHR site mutated like *in vivo* GFP reporter experiment |
| Probe synthesis | *sele* wild-type | GTAATGCCCCCTGTGACCCATATTGTCTCGCTCTTTCCTTTATA | TATAAAGGAAAGAGCGAGACAATATGGGTCACAGGGGGCATTAC | |
| Probe synthesis | *sele* mutation | GTAATGCCCCCTGattaaCATATTGTCTCGCTCTTTCCTTTATA | TATAAAGGAAAGAGCGAGACAATATGttaatCAGGGGGCATTAC | NHR site mutated like *in vivo* GFP reporter experiment |

Table shows primers used for cloning mouse *Nr2f2* into the pGEX2TK vector and DNA probes from the zebrafish *mrc1a* and *sele* enhancers.
